# Population dynamics of Arctic phytoplankton and mycoplankton reveal chytrid-mediated diatom bloom termination

**DOI:** 10.64898/2026.07.07.736921

**Authors:** Pauline C. Thomé, Ellen Oldenburg, Cora Hörstmann, Jürgen F. H. Strassert

**Affiliations:** Department of Environmental Microbiomics, Technische Universität Berlin, Berlin, Germany; Institute of Quantitative and Theoretical Biology, Heinrich Heine University Düsseldorf, Düsseldorf, Germany; Cluster of Excellence on Plant Sciences, Heinrich Heine University Düsseldorf, Düsseldorf, Germany; Deep-Sea Ecology and Technology, Alfred Wegener Institute Helmholtz Centre for Polar and Marine Research, Bremerhaven, Germany; Ecological Chemistry, Alfred Wegener Institute Helmholtz Centre for Polar and Marine Research, Bremerhaven, Germany

**Keywords:** Chytridiomycota, Fungi, Parasitism, Arctic Ocean, Sea ice, MOSAiC, *Tara*

## Abstract

Chytrids are unicellular fungi that infect and degrade phytoplankton as parasites or saprotrophs. They impact not only food availability and quality in surface waters but also carbon cycling and sequestration. So far, their ecological significance has mostly been investigated for freshwater environments, whereas observations for marine environments are scarce — even though chytrids can be highly abundant there, too (as shown for the Arctic Ocean). To test the chytrids’ potential to control phytoplankton dynamics in the Arctic Ocean, we analysed metabarcoding and photosynthetic pigment data from two expeditions, *Tara* Polar Circle and MOSAiC; the latter providing a dense sampling transect across one year from the under-ice water column and sea ice samples. The phytoplankton communities of both environments were dominated by diatoms, with strong seasonal effects indicating blooms in the water column. Chytrids dominated fungal communities in both environments and revealed a strong cryo–pelagic coupling. They were especially abundant during the sea ice melt in water samples and in ice-associated (sympagic) samples, where they represented >2% and up to 61%, respectively, of all combined reads assigned to chytrids or phytoplankton. Co-occurrences of the two most abundant chytrid taxa with some of the most abundant diatom taxa and niche differentiation from other potential diatom parasites are consistent with the chytrids’ critical role in controlling diatom blooms, especially in sympagic habitats.

## Introduction

Unicellular fungi belonging to chytrids (Chytridiomycota) are assumed to be of global importance as regulators of phytoplankton populations. While many chytrids are found to parasitize phytoplankton, including, for example, diatoms, dinoflagellates and cyanobacteria (Frenken et al., 2017; Thomé et al., 2024; Van den Wyngaert et al., 2022), others live as saprotrophs or intermediate forms that degrade dead or moribund phytoplankton biomass. Their flagellated zoosporic stage proliferates by feeding on the cells while growing into sporangia, in which new zoospores are built (Ibelings et al., 2004). Chytrids can thereby accelerate bloom termination (Frenken et al., 2016), make carbohydrates and nutrients more accessible to herbivores by host fragmentation (Frenken et al., 2020, 2017) and constitute a valuable food source for zooplankton that are enriched in fatty acids and sterols (Gerphagnon et al., 2019). Moreover, chytrids contribute to retaining hosts that can form fast-sinking colonies or substrates in surface waters, potentially impacting the global carbon cycle and organic matter sequestration (Boetius et al., 2013; Kagami et al., 2007; Riebesell et al., 1991). Our incipient knowledge about phytoplankton–chytrid interactions has largely been derived from freshwater systems, while chytrids remained unrecognised in marine environments until recently. In 2014, the first parasitic marine chytrid (infecting a dinoflagellate) was described by Lepelletier et al. (2014).

Subsequent metabarcoding studies gave first insights into the chytrids’ global occurrence. They were found in various temperate and polar marine ecosystems being particularly abundant in the Arctic, where they could represent >90% of the whole fungal community (Comeau et al., 2016; Hassett and Gradinger, 2016; Kilias et al., 2020). High abundances were specifically found in and under the sea ice (SI; Comeau et al., 2011; Kilias et al., 2020), and it has been speculated that, together with other fungi, chytrids may even substitute algae as the base of the Arctic marine food web (Hassett et al., 2017).

Understanding the chytrids’ potential to control Arctic phytoplankton populations is of growing interest as climate-change-induced SI loss and elevated irradiance have caused an increase in pelagic primary production of 30% already between 1998 and 2012 (Arrigo and Van Dijken, 2015), and until the end of the century an increase of 550% in ice-algae productivity is expected in the central Arctic (Tedesco et al., 2019). Yet, due to the limited number of samples that are usually analysed and the chytrids’ quickly fluctuating abundances, meaningful discoveries concerning their dynamics remain highly sporadic, providing only a fragmented understanding of their role in controlling phytoplankton dynamics. Furthermore, chytrids are commonly reported solely in relation to the fungal community (e.g. Comeau et al., 2016; Hassett, 2020; Hassett and Gradinger, 2016; Jeffries et al., 2016; Richards et al., 2015) and reference databases are usually extremely biased towards Dikarya (Carradec et al., 2018; Morales et al., 2019) with only a few chytrid references to allow correct mapping, often leaving many fungal sequences unassigned (Hassett et al., 2019). Thus, a comprehensive picture of the Arctic chytrids’ seasonality, habitat and host association is still lacking.

Thanks to the MOSAiC expedition, the largest-scale Arctic research expedition to date (Mock et al., 2022), data from a unique sampling transect across the central Arctic Ocean are now available, which allow us to establish a comprehensive understanding of Arctic marine microbial life throughout the year. Using metabarcoding data from this expedition, as well as the *Tara* Polar Circle expedition (Alberti et al., 2017), we aimed to provide a context for the chytrids’ episodically high abundances in the Arctic marine environment. We quantified chytrid abundances and linked them to i) habitat by comparing pelagic and sympagic (sea ice) communities, ii) seasonality by taking advantage of the dense sampling transect, iii) environmental conditions by including temperature, salinity and pigment concentrations and iv) phytoplankton host association by relating chytrids to the phytoplankton community and identifying putative host–parasite pairs. We provide new insights into the role of chytrids at the base of Arctic marine food webs by revealing chytrid associations with the sympagic habitat and with the season of SI melt, as well as with single diatom hosts rather than with total diatom biomass.

## Materials and methods

### Metabarcoding and environmental data processing

Metabarcoding data was analysed from both the V4 region and the V9 region of the 18S rRNA gene sequences generated during the MOSAiC campaign (2019–2020; V9, Chamberlain and Bowman, 2022; V4, Metfies et al., in revision) and the *Tara* Polar Circle campaign (2013; NCBI BioProject PRJEB9737, note that some V9 samples are mislabeled as V4; Alberti et al., 2017). Marker regions were amplified with the primer pairs 528iF/964iR (MOSAiC, V4; Metfies et al. 2020), 1380F/1510R (MOSAiC, V9; Amaral-Zettler et al., 2009) and 1389F/1510R (*Tara*, V9; Amaral-Zettler et al., 2009). Other sampling and sequencing procedures were described elsewhere for both data sets (Vargas et al., 2015; Alberti et al., 2017; Chamberlain et al., 2025; Metfies et al., in revision). Among these data, we distinguished between under-ice water (UIW) samples and sea ice (SI) samples from the ice-covered central Arctic Ocean (MOSAiC), and ice-free water (IFW) samples from the open Arctic Ocean marginal seas (*Tara* Polar Circle). Primers were removed with cutadapt v4.8 using default settings and -O 6. Reads were filtered (maxEE = 2), trimmed, dereplicated, denoised and merged, and chimeras were removed with DADA2 v1.36.0 in R v4.5.0 using RStudio v2024.12.1 with default settings except the specified flag. The amplicon sequence variants’ (ASVs) taxonomy was assigned using the PR^2^ database v5.0.0 (bootstrap threshold 50; Guillou et al., 2013). Publicly available sequences of 44 not yet included chytrid taxa were added to the database to roughly represent all lineages on a family level (Seto et al., 2023; Thomé et al., 2024; Van den Wyngaert et al., 2022).

Furthermore, 18S rRNA gene sequences of 38 previously not included potential non-chytrid diatom parasites were added: members of the novel chytrid-like clade 1 (NCLC1), aphelids, rozellids, oomycotes, Pirsoniales, *Cryothecomonas*, *Solenicola* and *Phagomyxa* (e.g. Chambouvet et al., 2019; Prokina et al., 2024; Seto et al., 2023). Mislabeled chytrids in PR^2^ (Olpidiomycota) were excluded.

Environmental data for the pelagic were collected for the pigments chlorophyll *a*, chlorophyll *c* and fucoxanthin (Hoppe et al., 2023a, 2023b; van Leeuwe et al., 2023), and for the oceanographic parameters salinity and temperature (Schulz et al., 2023). Pigment concentrations, which were occasionally measured at different times and places than the DNA was sampled, were interpolated using the functions *Tps()* and *predict()* from the R package *fields* v17.1. As a proxy for blooms in the water column, pigment anomalies were computed as pigment concentration minus its standard deviation (Tedesco and Vichi, 2014), and positive values were considered to represent blooms.

Temperature and salinity were measured more densely and deviations in time and space from DNA sampling points were taken into account by linear interpolation between oceanographic measurements.

### Community analysis

To disentangle phytoplankton and mycoplankton dynamics, unicellular eukaryotic lineages were grouped as “protists” (including unicellular fungi) and analysed. This selection served as reference for calculating phytoplankton proportion. Replicates from the same date, location and depth, as well as from the same sampling point (different size fractions) were merged. Eukaryotic phytoplankton were selected by including photoautotrophic lineages within the Ochrophyta, Dinophyta (and related taxa; after Schneider et al. (2020), Hörstmann et al. (2022) and Bruhn et al. (2024)), Haptophyta, Archaeplastida and Cryptophyta. Bacillariophyta is here referred to as diatoms. For the selection of the most abundant phytoplankton, the 30 most abundant taxa among all analysed samples were combined with the three most abundant taxa per sample. Prokaryotic phytoplankton were retrieved for additional verification by selecting the phylum Cyanobacteria from 16S rRNA gene data (MOSAiC; Chamberlain and Bowman, 2022).

The community data was analysed with the R packages *microViz* v0.12.7 and *phyloseq* v1.52.0 (Barnett et al., 2021; McMurdie and Holmes, 2013). Samples comprising less than 1,000 reads of protist taxa were removed from the analyses. Two datasets were selected from the pelagic samples: i) a full dataset including all samples up to 115 m water depth, which was the maximum depth at which chlorophyll *a* (Chl *a*) concentrations reached ≥0.12 µg L^−1^ (defined as bloom threshold) and ii) a time series including one sample from each day with the highest Chl *a*. For SI, all available samples were used; each sample corresponded to a 10 cm section of an ice core (total depth up to 2.7 m). Pearson correlations from Sparse Partial Least Squares regressions between taxa abundances were computed from log-ratio-transformed data with the *spls()* and *cim()* functions from the *mixOmics* v6.32.0 R package (Lê Cao et al., 2011). To account for the compositionality of metabarcoding data, proportionality was additionally assessed with the *propr* function from the *propr* v5.1.6 R package (Quinn et al., 2017) using a false-discovery-rate threshold of 0.05. Co-variations in proportional abundance over time between putative host–parasite pairs were visualised by *z*-scores, i.e. number of standard deviations from the mean of abundances. The map was created with the R packages *sf* v1.0.21 and *rnaturalearth* v1.1.0.

## Results

To investigate the chytrids’ impact on Arctic phytoplankton blooms, we analysed 18S V4 and V9 metabarcoding amplicons obtained from under-ice water (UIW) collected on 217 sampling days (October 2019–October 2020), from sea ice (SI) collected on 13 sampling days (November 2019–October 2020) and from ice-free water (IFW) collected on 23 sampling days (May–November 2013; Fig. 1, Table S1). Especially the V9 data represents a dense UIW sampling transect following the transpolar drift from north of the Laptev Sea across the central Arctic Ocean to the ice edge in the Fram Strait, followed by a transit back to the centre of the ice cover near the North Pole (Fig. 1; Fong et al., 2024). The total read numbers varied between 2,090 and 7,757,673 across all datasets after merging replicates and selecting reads assigned to protists (see Materials and methods). Due to substantial variation in sequencing depth across samples, read counts were not rarefied or otherwise normalised. Instead, all downstream analyses were based on relative (compositional) abundances, ensuring that differences in sequencing depth do not bias patterns of community composition. The results should therefore be interpreted in a relative rather than absolute sense. The V9 samples from the ice-covered Arctic (MOSAiC) generally had lower sequencing depths than the V4 samples (Fig. S1, Table S1). However, comparison of the results inferred from the two markers in UIW samples showed a largely coherent protist community composition on those dates and water depths from which both markers were sequenced (Fig. S2). To take advantage of the central Arctic Ocean year-round sampling, subsequent results are based on the 18S V9 data.

**Fig. 1:**
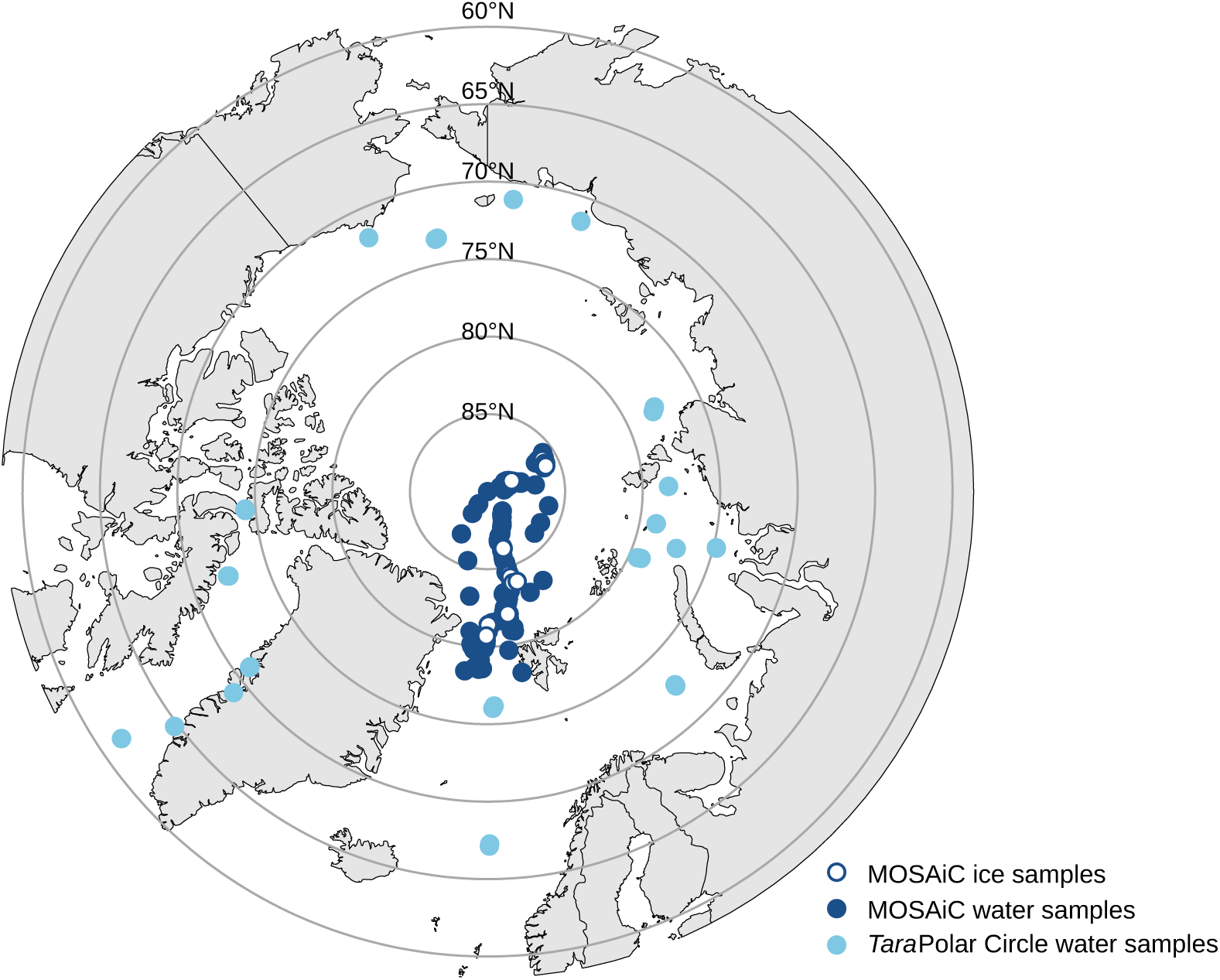
Sampling sites in the ice-covered Arctic Ocean (MOSAiC campaign) and ice-free Arctic Ocean (*Tara* Polar Circle campaign). Under-ice water (dark blue), ice-free water (light blue) and sea-ice (white) samples are distinguished.

In the ice-covered central Arctic Ocean, UIW phytoplankton peaked in spring and summer (as well as in November), as shown by increasing chlorophyll *a* (Chl *a*) concentrations and increasing numbers of phytoplankton reads in proportion to all protist reads (Fig. 2a, b), which correlated closely (Pearson’s *r* = 0.63, *p* < 0.005; Fig. S3a). The proportion of protists that was assigned to phytoplankton in single samples ranged between 3.5% in February and 84% in November (Fig. 2b); the latter likely reflecting the ability of diatoms to survive under extremely low light levels rather than an active bloom (Berge et al., 2015; Joli et al., 2024). Positive Chl *a* anomalies, calculated as pigment concentration minus its standard deviation (Tedesco and Vichi, 2014), were found to set in at the end of May (Fig. 2b), indicating the beginning of the UIW spring bloom as estimated by Hoppe et al. (2024). Diatom blooms were also well described by changes in Chl *a*, which showed a stronger correlation with their abundances than the diatoms’ additional pigments fucoxanthin and chlorophyll *c* (Pearson’s *r* = 0.36, *p* < 0.005; Fig. S3b–d). The latter two pigments showed positive anomalies only later in spring (mid-June; Fig. S4).

**Fig. 2:**
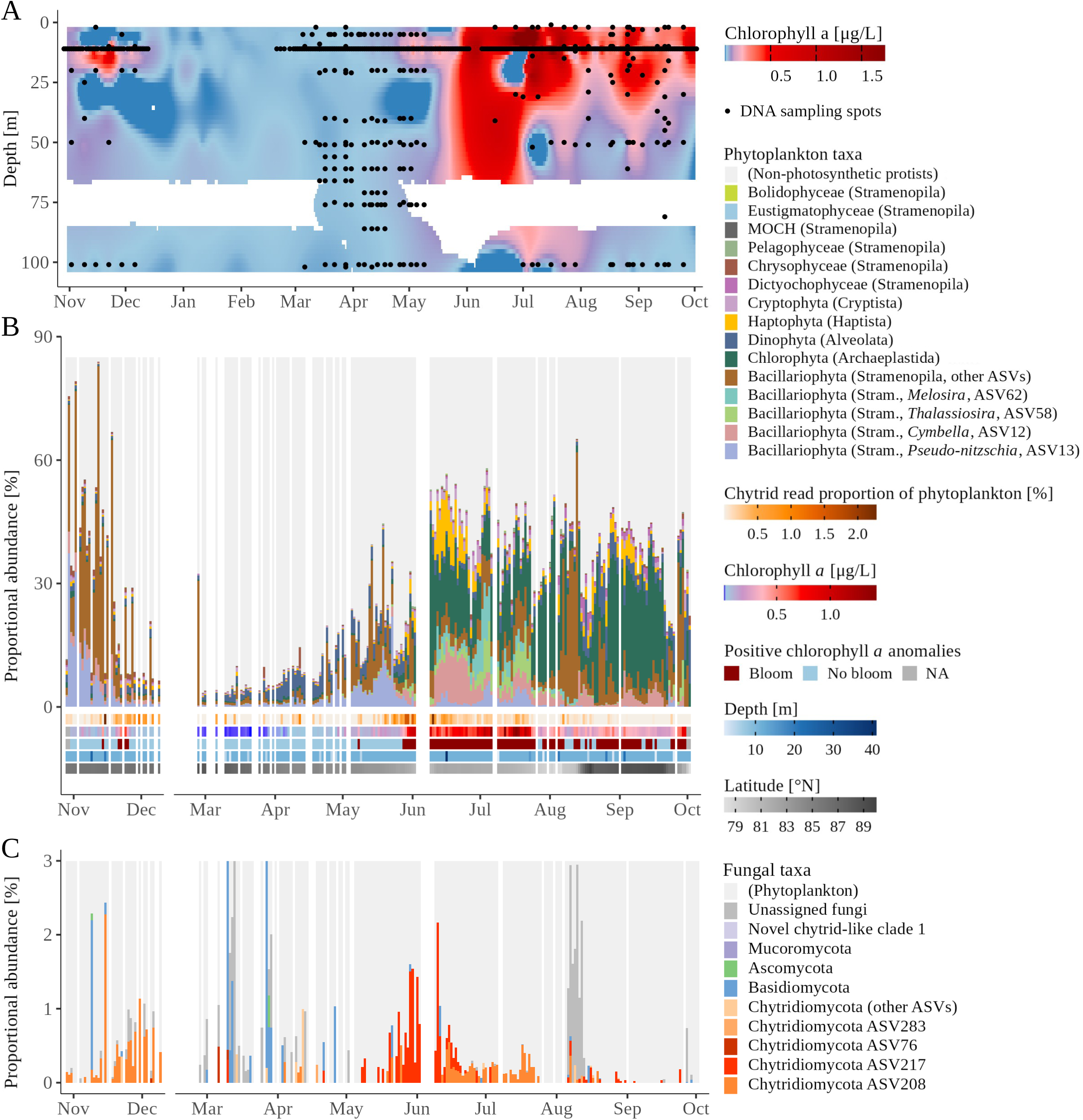
Environmental and community data in the under-ice water. (A) Interpolation of chlorophyll *a* concentrations (Chl *a*; colour gradient) and sampling points; (B) proportion of phytoplankton among protists (see Methods) and proportion of chytrids within the chytrid + phytoplankton communities, Chl *a* anomaly, depth of the time-series samples and latitude (note the northwards transition in August), time-series data; (C) proportion of fungi within the fungi + phytoplankton communities, time-series data. Note that the y-axis is cropped for better visualisation of chytrids; to see data exceeding the displayed range, please refer to Fig. S16. White space: no data.

### Diatoms dominate Arctic Ocean phytoplankton communities

Phytoplankton communities in UIW were overall dominated by diatoms (45% of total phytoplankton; Fig. S5a). This was particularly true for spring and November (Fig. 2b). Here, *Pseudo-nitzschia* represented the most abundant phytoplankton (ASV13, up to 62% in single samples), when Chl *a* was elevated compared to winter levels but after and before it exceeded bloom thresholds (Fig. S6). *Cymbella* represented the most abundant diatom during the June bloom (ASV12, up to 27% in single samples), followed by *Thalassiosira* (ASV58, up to 23% in single samples) and *Melosira* (ASV62, up to 33% in single samples) in July, *Chaetoceros* (ASV48, up to 83% in single samples) in early August and *Actinocyclus* (ASV45, up to 47% in single samples) in late August (Fig. S6). Chlorophyta increased in proportion starting in June and remained abundant until autumn, making up almost a quarter of all phytoplankton reads (Figs. 2b, S5a). Here, *Micromonas* was particularly abundant, vastly dominating blooms towards the end (ASV4, up to 88% in single samples; Fig. S6).

Photoautotrophic dinophytes represented about 17% of total phytoplankton and were most abundant between December and May (Figs. 2b, S5a), with *Lepidodinium* dominating the phytoplankton communities initially after the return of light in March and April (ASV39, up to 53% in single samples; Fig. S6). Other lineages accounting for at least 1% of the phytoplankton community belonged to Haptophyta (5.6% of total phytoplankton), with *Phaeocystis* (ASV47, up to 24% in single samples) and *Chrysochromulina* (ASV135, up to 19% in single samples) being most abundant in June and late August/early September, respectively (Fig. S6), Cryptophyta (4.2%), Chrysophyceae (2.5%) and Dictyochophyceae (1.7%; Fig. S5a). In IFW, phytoplankton communities were also overall dominated by diatoms (50% of total phytoplankton; Fig. S5b). They dominated the phytoplankton throughout most of the summer and were complemented mostly by dinophytes (>70% in single samples; Fig. S7a). In contrast to UIW, chlorophytes accounted for only low abundances in IFW (4% of total phytoplankton; Fig. S5b).

In SI, the overwhelming dominance of diatoms was even more pronounced (58% of all phytoplankton, Figs. 3a, S5c). The most abundant diatom ASVs that could be taxonomically assigned to the genus level were affiliated to *Sellaphora*, *Pseudo-nitzschia*, *Navicula* and *Pleurosigma* (Fig. S8). Diatoms were abundant throughout all ice depths, while a spatial structuring was found for dinophytes, which were located mainly in the middle and lower parts of the ice cores (14.2% of total SI phytoplankton; especially *Polarella* and *Lepidodinium*), and in chrysophytes, which were mainly located in the upper parts (12.7%; Fig. 3a). Further abundant taxa belonged to the Chlorophyta (11%) and Cryptophyta (1.8%; Fig. S5c). Compared to the UIW communities, a notably high number of ASVs remained unassigned in SI beyond taxonomic class level (Fig. S8).

**Fig. 3:**
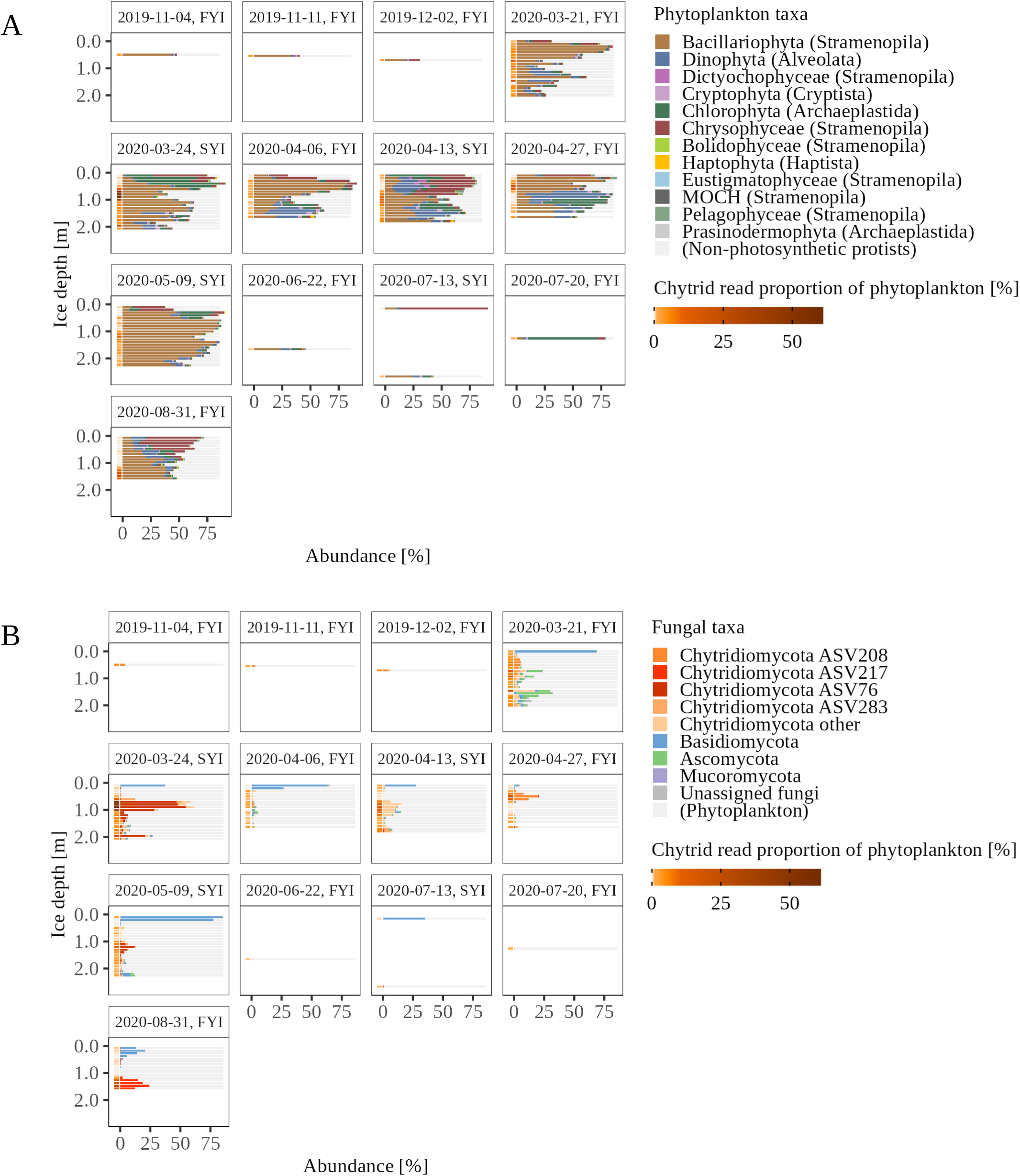
Phytoplankton and mycoplankton communities in the sea ice. (A) Proportion of phytoplankton among protists and proportion of chytrids within the chytrid + phytoplankton communities; (B) proportion of fungi within the fungi + phytoplankton communities. FYI: first-year sea ice; SYI: second-year sea ice. White space: no data.

### Chytrids are the most abundant Arctic marine fungi

Chytrids vastly dominated mycoplankton communities, in particular in the ice-covered Arctic, comprising 69% (UIW), 65% (SI) and 17% (IFW; Fig. S9) of all fungal reads. They were mainly comprised of ASV208 and ASV217 in UIW, where they oscillated throughout the year (Figs. 2c, 3b), and were more diverse in SI. That is, 13 ASVs accounted for 90% of all chytrid sequences in SI but only two ASVs in UIW. In proportion to phytoplankton (i.e. chytrid proportion of the combined chytrid + phytoplankton communities), chytrids reached up to 61% in single sympagic samples (i.e. in SI), which is significantly higher than in UIW and IFW (up to 2.3% and 2.2%, respectively; Fig. 4). In UIW, the relative abundance of chytrids was elevated during blooms (Fig. S10a). This is in contrast to IFW, where higher chytrid proportions were found at greater depths, with a maximum of 7.8% in one sample obtained from 651 m depth in the Norwegian Sea (Figs. S10b, S11). Members of the Basidiomycota, accounting for 9% of the total mycoplankton, were the only other assigned fungi that represented >1% across all UIW time-series samples (22% of the sequences remained unassigned; Fig. S9a), and while chytrids often represented the only fungi in single samples, Basidiomycota occasionally replaced them (Figs. 2c, S16). The mycoplankton in SI and IFW were more diverse, with Ascomycota (SI: 11%, IFW: 6%), Zoopagomycota (IFW: 3.2%) and Basidiomycota (SI: 22%, IFW: 1.7%) each reaching abundances of at least 1% across all time-series samples, while 72% of the reads remained unassigned in IFW and 3% remained unassigned in SI (Figs. 3b, 7b, S9b, c). Across the SI cores, Basidiomycota were usually found in the uppermost parts near the surface (*Sporobolomyces* and *Mrakia* dominated here) and chytrids were found mostly in the middle and lower sections near the pelagic (Fig. 3b).

**Fig. 4:**
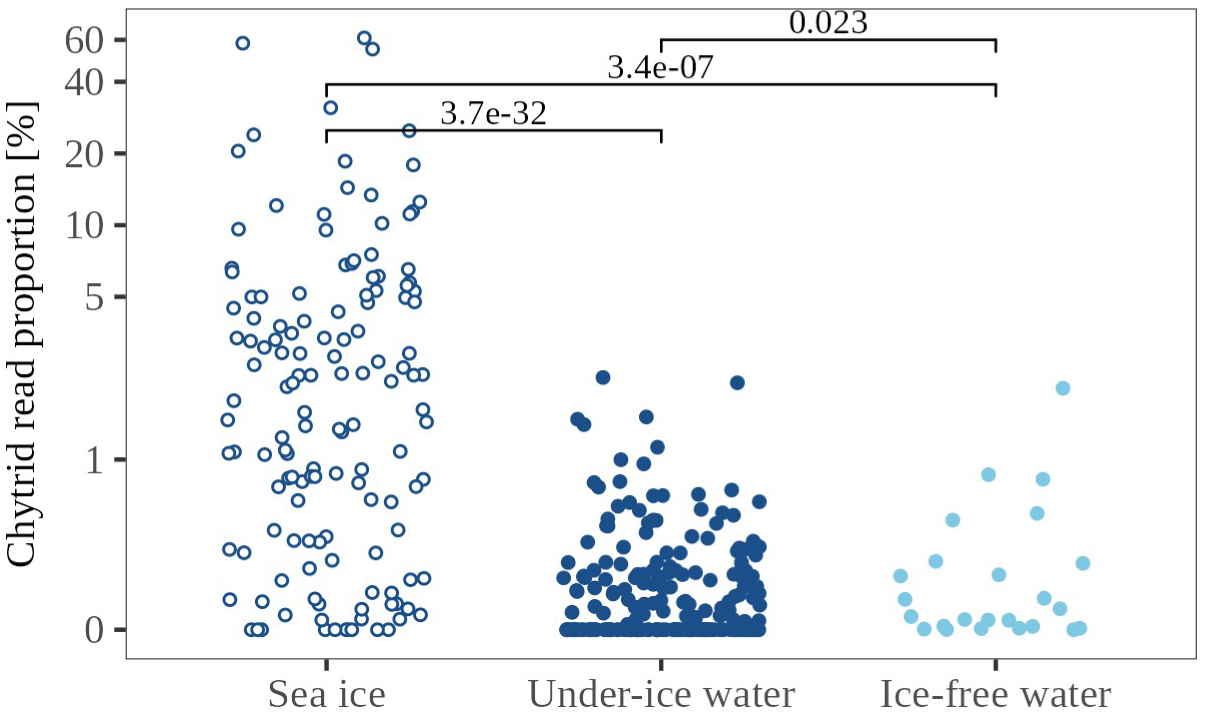
Proportion of chytrids within the chytrid + phytoplankton communities in the different environments. Means were compared with the Wilcoxon test and adjusted *p*-values are reported. Under-ice water and ice-free water: time-series data. Note the log-transformed scale.

### Arctic chytrids respond to diatom blooms and show niche differentiation from other diatom parasites

The same two chytrid ASVs (208 and 217) were abundant in SI and UIW. In a phylogenetic analysis, they formed a clade with other environmental chytrid sequences, which fell within the Lobulomycetales and which were closely related to diatom parasites, including *Zygorhizidium affluens* (Beakes et al., 1988; Van den Wyngaert et al., 2022; Fig. S12). In UIW, chytrids were most abundant from May to end of July as well as in November, i.e. when diatoms were most abundant, too (Fig. 2b). We found seasonal dynamics in the time series between chytrid ASV208 and ASV217 and some of the most abundant phytoplankton ASVs (Fig. 5). *Pseudo-nitzschia* (ASV13) and the haptophyte *Phaeocystis* (ASV47) correlated significantly with chytrid ASV208, and the diatoms *Thalassiosira* (ASV58), *Melosira* (ASV62) and *Cymbella* (ASV12) showed additional proportionality (Fig. 5). These pairs’ covariations were based on simultaneous peaks between May and July when only one other potential diatom parasite (*Cryothecomonas* ASV33) was abundant (Fig. 6a, b). The chytrids’ dominance among the diatom parasites was superseded from late June by *Pirsonia* (Stramenopila) and taxa assigned to the Labyrinthulomycetes (Sagenista) and diverse *Cryothecomonas* (Cercozoa). From late August to October, fungi and thereby chytrid populations decreased to negligible abundances, coinciding with a pronounced dominance of *Micromonas* (chlorophyte) in the phytoplankton community and *Pirsonia*, Labyrinthulomycetes and *Cryothecomonas* in the parasite community (Figs. 2c, 6a, b, S6). In SI, the same chytrid ASV208 and abundant phytoplankton correlated, including *Pseudo-nitzschia* (ASV13) and *Cymbella* (ASV12). No niche differentiation among the diatom parasites was found, with the exception of the first-year sea ice core from March 21, where *Cryothecomonas* dominated in the bottom ice, matching the proportions in the water below (Fig. S13). Other potential diatom parasites such as oomycotes, aphelids, rozellids or species belonging to the opisthosporidian NCLC1 clade (Chambouvet et al., 2019; Garvetto et al., 2018; Käse et al., 2021; Tillmann et al., 1999) were almost absent in the data.

**Fig. 5:**
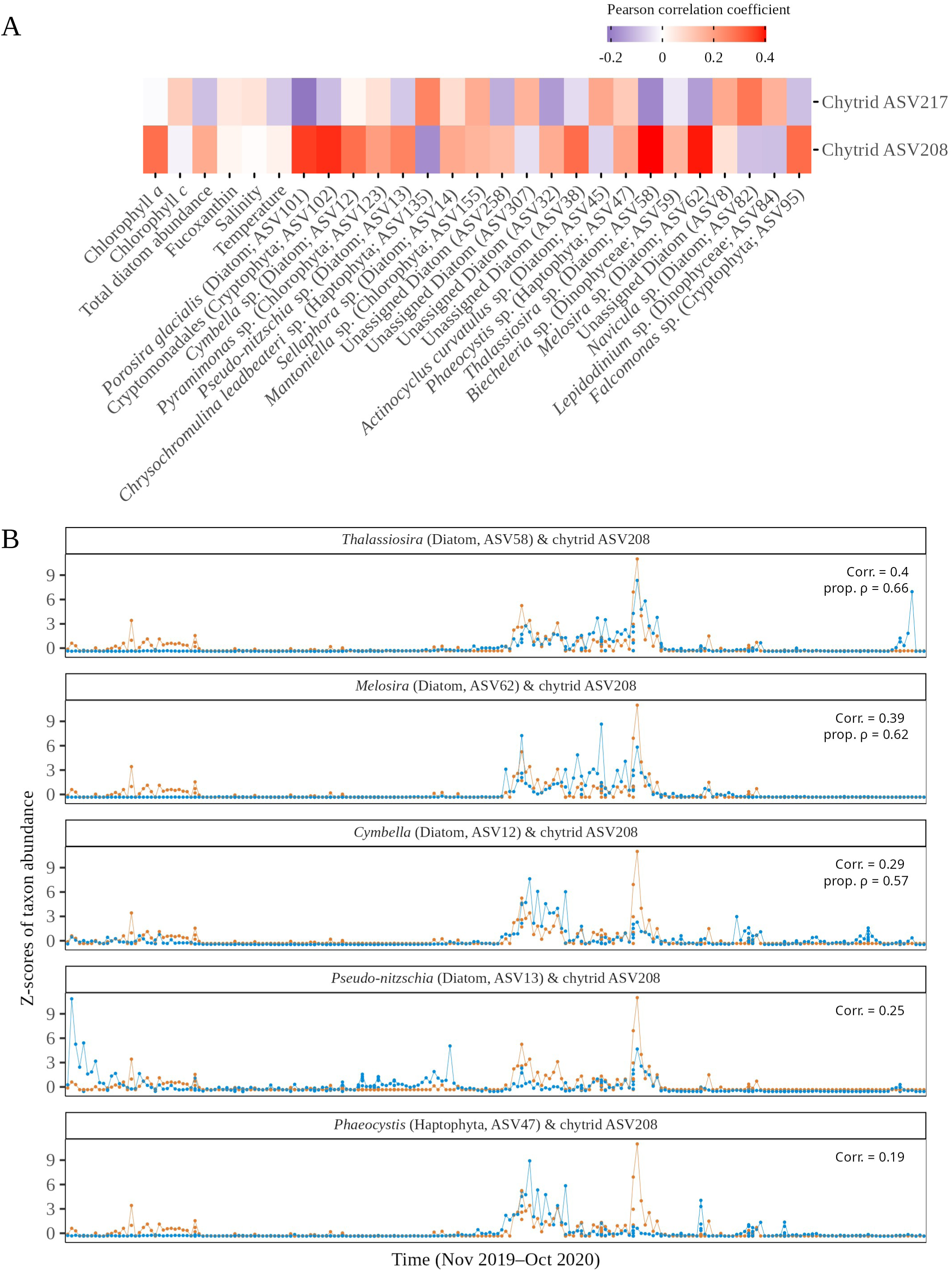
Correlations between individual chytrids and the most abundant phytoplankton ASVs as well as selected environmental parameters in the under-ice water, full dataset. (A) Pearson correlations from Sparse Partial Least Squares regressions from the full dataset. Only phytoplankton ASVs with correlation coefficients >0.1 with at least one chytrid ASV are considered; (B) *z*-scores of raw read counts (taxon abundance), Pearson correlation coefficients and proportionality coefficient *rho* (shown only if significant) of phytoplankton ASVs and of the chytrid ASV208 (shown as example). Orange: chytrid; blue: phytoplankton.

**Fig. 6:**
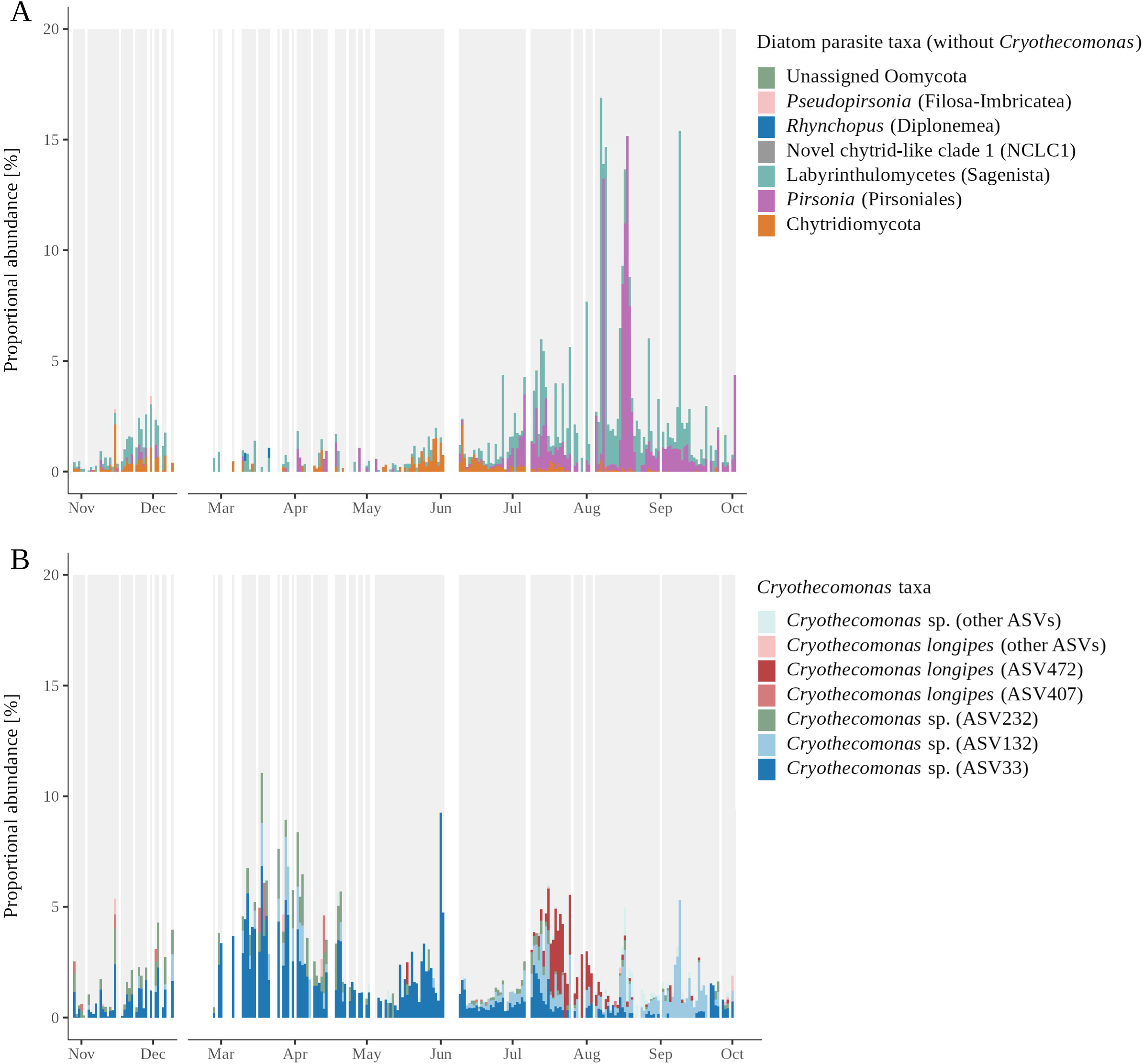
Proportion of potential non-chytrid diatom parasites within the phytoplankton + parasite communities, time-series data. (A) Non-*Cryothecomonas* lineages (debated diatom parasites such as Syndiniales were not included), grey bars: phytoplankton + *Cryothecomonas*; (B) *Cryothecomonas* lineages, grey bars: phytoplankton + non-*Cryothecomonas.* Note, ASV33 parasitizes on a diatom species (*Guinardia delicatula*) that was not present in our data (see Discussion for more information).

## Discussion

To reveal the impact of chytrids on Arctic phytoplankton dynamics, we analysed metabarcoding datasets generated during the MOSAiC and *Tara* Polar Circle campaigns. We focused on changes across samples to minimise potential errors caused, for example, by varying rRNA gene copy numbers in diverse lineages (Gong and Marchetti, 2019; Lofgren et al., 2019). Our study provides a framework for understanding the seasonality of the highly dynamic chytrid abundances in the Arctic Ocean. The results allow us to attribute the emerging evidence of their seasonal importance in pelagic phytoplankton–chytrid communities to the transition from spring to summer during the sea ice (SI) melt (May–July; note, spatial differences were not investigated). The rising chytrid abundance in under-ice water (UIW) in May coincided with both the increase in pelagic chlorophyll *a* (Chl *a*) concentrations and the increase in phytoplankton reads up to bloom levels. Diatoms dominated the phytoplankton community during this time, and our data strongly suggest that the termination of the diatom spring bloom was mediated by chytrids.

The strong cryo–pelagic coupling of chytrid taxa uncovered here provides the first evidence for the importance of SI as a reservoir for chytrids that seeds the pelagic below — a phenomenon that was repeatedly hypothesised for both fungi and algae but has not yet been confirmed (Hassett and Gradinger, 2016; Ardyna et al., 2020, Kilias et al., 2020). Chytrids that were abundant in SI and UIW belonged to the same ASVs. Corroboration for the cryo–pelagic coupling is further given by the timing of the UIW chytrid population increase, which fell together with the seasonal onset of bottom-ice melt (Raphael et al., 2024; this study), as well as by concordantly abundant algal ASVs in both SI and UIW (e.g. *Navicula*, *Pseudo-nitzschia*, *Melosira* and *Polarella glacialis*).

The cross-season dominance of chytrids in both SI and UIW fungal communities is in line with other reports of high chytrid abundances sporadically found in the ice-covered Arctic Ocean (Hassett et al., 2017; Hassett and Gradinger, 2016; Kilias et al., 2020). Interestingly, comparable proportions have not yet been documented for any other habitat worldwide. In non-Arctic marine environments, including the Southern Ocean, chytrids usually occur in low abundances compared to Dikarya (Duan et al., 2018; Morales et al., 2019; Pham et al., 2021; Taylor and Cunliffe, 2016; Tisthammer et al., 2016), and in freshwater, where they can reach high infection prevalence in phytoplankton (Frenken et al., 2016), chytrid dominance among the fungal community is observed only periodically (Van den Wyngaert et al., 2022). It is remarkable in this context that exceptions are known especially from environments with varying salinity like brackish benthos (Ilicic et al., 2024) or coastal regions with river input (Comeau et al., 2016; Debeljak and Baltar, 2023; Gutiérrez et al., 2016; Taylor and Cunliffe, 2016). Hence, chytrids seem to tolerate salinity variations as present in the SI brine channels, making them an ideal environment in which they can easily encounter their hosts during diatom blooms. But even though chytrids reach proportions of up to two-thirds in Arctic SI chytrid + phytoplankton communities, so far, they could not be linked to phytoplankton dynamics.

Since chytrids comprise parasitic and saprotrophic lineages (and even intermediate forms that have the capacity to use both detritus and host cells to complete their life cycle; Frenken et al., 2017), deducing their lifestyle based on metabarcoding data is challenging. Although fluorescence *in situ* hybridisation or single-cell sequencing approaches could not be applied here, the following observations argue strongly for parasitic interactions between several chytrid and diatom lineages: i) chytrid peak abundances were mostly caused by single ASVs (typical for host-specific and density-dependent parasites; Taylor and Cunliffe, 2016); ii) chytrid abundances within the chytrid + phytoplankton community were much more strongly related to single diatom ASVs than to the total diatom community or to Chl *a*; iii) chytrids correlated with diatom lineages that are well-known chytrid hosts (*Pseudo-nitzschia* (Hanic et al., 2009), *Thalassiosira* (Gutiérrez et al., 2016), *Navicula* (Scholz, 2014) and *Melosira* (Gleason et al., 2014)); iv) chytrids clustered phylogenetically with the diatom parasites *Zygorhizidium affluens* and isolate Fragilaria-MDA25 (however, this needs to be considered with caution as multiple lifestyle transitions are known across the chytrid phylum; Thomé et al., 2024); v) absence of other potential diatom parasites during chytrid peak abundances. As numerous chytrid species were shown to infect highly diverse hosts (Kagami et al., 2021; Lepelletier et al., 2014; Pou-Solà et al., 2026; Van den Wyngaert et al., 2022), the varying correlation strength between host–parasite pairs could be interpreted as an indication of generalism. It is noteworthy in this context that the haptophyte *Phaeocystis* (showing a weak correlation) has not been observed as a chytrid host before. To corroborate the host–parasite pairs presented here, and to rule out spurious correlations, we also included other potential, non-chytrid diatom parasites in our analyses as well as zooplankton as potential predators of both phytoplankton and chytrids. Correlations were neither found between the parasitic chytrids and zooplankton nor between the putative phytoplankton hosts and zooplankton (data not shown). Furthermore, based on the observed niche differentiation of diatom parasites throughout the year, the impact of non-chytrid parasites on diatom dynamics during the Arctic spring bloom can be neglected. Only ASV33 co-occurred with chytrids during this period (Fig. 6a, b). The sequence of this ASV, however, was virtually identical to and clustered phylogenetically with the sequence of *Cryothecomonas aestivalis*, a species that parasitizes exclusively on a diatom that was not present in the samples and therefore did not affect our data (*Guinardia delicatula*; Drebes et al., 1996). Non-chytrid parasites are likely to regulate Arctic diatom dynamics over different time intervals than chytrids (e.g. *Cryothecomonas*; Dilliplaine et al., 2025) or infect other Arctic phytoplankton (e.g. MALVs or some *Cryothecomonas* lineages; Fig. S14; Egge et al., 2025). The impact that cyanobacteria can have on chytrid dynamics remains largely hidden here. The dynamics of some potential cyanobacterial host lineages were investigated, but they peaked at different times of the year than the diatom hosts. Statistical cross-comparisons between 16S and 18S rRNA gene amplicon datasets are not advisable, as differences in primer specificity, amplification efficiency and rRNA gene copy number variation between prokaryotic and eukaryotic organisms preclude quantitative inference across these fundamentally distinct marker systems (Větrovský and Baldrian, 2013).

The high abundances of chytrids in SI and UIW suggest they may serve as food for zooplankton, constituting an important link in the Arctic marine food web. The existence of such a link, postulated in the “mycoloop” hypothesis (Kagami et al., 2014), is supported by reports of disproportionally high consumption of chytrids by *Calanus* individuals in Arctic fjords during spring blooms (Cleary et al., 2017) as well as accumulation of host fatty acids and synthesis of sterols in diatom-infecting chytrids (Gerphagnon et al., 2019). The increase in chytrid abundance in spring matches the time when SI algal biomass and productivity reach a multiple of those in the pelagic (Gosselin et al., 1997; Gradinger, 2009; Arrigo, 2017). While the large temporal and spatial distance between MOSAiC’s SI Chl *a* measurement points (Fig. S15) prevents an unambiguous linkage to the DNA data, we nonetheless know that the ice-algal spring bloom represents the first peak in nutritional quality of primary producers (Søreide et al., 2010). This peak is essential for zooplankton, which feed on ice algae and rely on high-quality food for reproduction after fatty acid depletion during the winter (Leu et al., 2011). As seasonal decoupling of zooplankton egg production and ice-algal spring blooms is ongoing (Søreide et al., 2010), chytrids may become more relevant by compensating for periods of low food quality.

In this study, we provide strong support for the chytrids’ role in controlling diatom dynamics. Under climate change scenarios, increasing irradiance — resulting in decreased ice thickness and snow cover as well as increased rainfall and melt pond formation (Lannuzel et al., 2020; Muilwijk et al., 2024) — is expected to enhance ice-algal primary productivity in the central Arctic by up to 6.5-fold in the future (Tedesco et al., 2019). These conditions may favour chytrid parasitism, as chytrid epidemics are not only host density-dependent (Ibelings et al., 2004) but also increase when their hosts are light-stressed (Hassett and Gradinger, 2016). The chytrids’ importance as an alternative food source for zooplankton may increase as well. Elevated irradiance and temperature were shown to lower the nutritional quality of SI algae (Leu et al., 2010) and to accelerate diatom bloom degradation by chytrids (Frenken et al., 2016), respectively. In the more distant future, in contrast, it is also possible that other phytoplankton lineages will dominate an ice-free Arctic Ocean, causing a shift in the parasite community away from chytrids. Our results may underline that the seasonal transition of peak productivity from the sympagic to the pelagic realm is shifting from early July to late May — this would be earlier than predicted under the RCP8.5 scenario for the second half of the century (Ardyna and Arrigo, 2020; Tedesco et al., 2019). The finely tuned timing of ice-algal bloom peaks with zooplankton maturation and egg production is already disturbed (Leu et al., 2011; Post, 2017; Søreide et al., 2010), stressing once more the importance of understanding these diverse food web links.

## Acknowledgments

We are grateful to Ovidiu Popa for his methodological advice. We further thank Clara J. M. Hoppe, Niels Fuchs, Maria Van Leeuwe and Emelia J. Chamberlain for their support with data access and curation, Danny Ionescu for helping with the pigment analysis, Kenneth Dumack for the phylogenetic assignment of *Cryothecomonas* ASVs, Michael T. Monaghan for valuable discussions and Mina Bizic for critical comments on the manuscript. Pauline C. Thomé acknowledges support from the federal Elsa Neumann Scholarship (NaFöG, Berlin) and Jürgen F. H. Strassert acknowledges support from the German Research Foundation (DFG; grant STR 1349/2-2, project no. 1-5003006-01-EF).

## Author contributions

Pauline C. Thomé: Conceptualisation, Funding acquisition, Data curation, Formal analysis, Investigation, Methodology, Project administration, Software, Visualisation, Writing — original draft, Writing — review & editing. Ellen Oldenburg: Methodology, Writing — review & editing. Cora Hörstmann: Methodology, Writing — review & editing. Jürgen F. H. Strassert: Conceptualisation, Funding acquisition, Investigation, Methodology, Project administration, Supervision, Writing — original draft, Writing — review & editing.

## Data availability statement

Data are available under NCBI BioProjects PRJEB9737 (*Tara* Polar Circle) and PRJNA895866 (MOSAiC 18S V9 and 16S). 18S V4 data are available upon request. Pigment data can be obtained from PANGAEA under DOIs 10.1594/PANGAEA.955763, 10.1594/PANGAEA.963277 and 10.1594/PANGAEA.962597, and hydrographic data can be obtained from the Arctic Data Center under DOI 10.18739/A21J9790B.

