## Supplementary figures for "Population dynamics of Arctic phytoplankton and mycoplankton reveal chytrid-mediated diatom bloom termination"

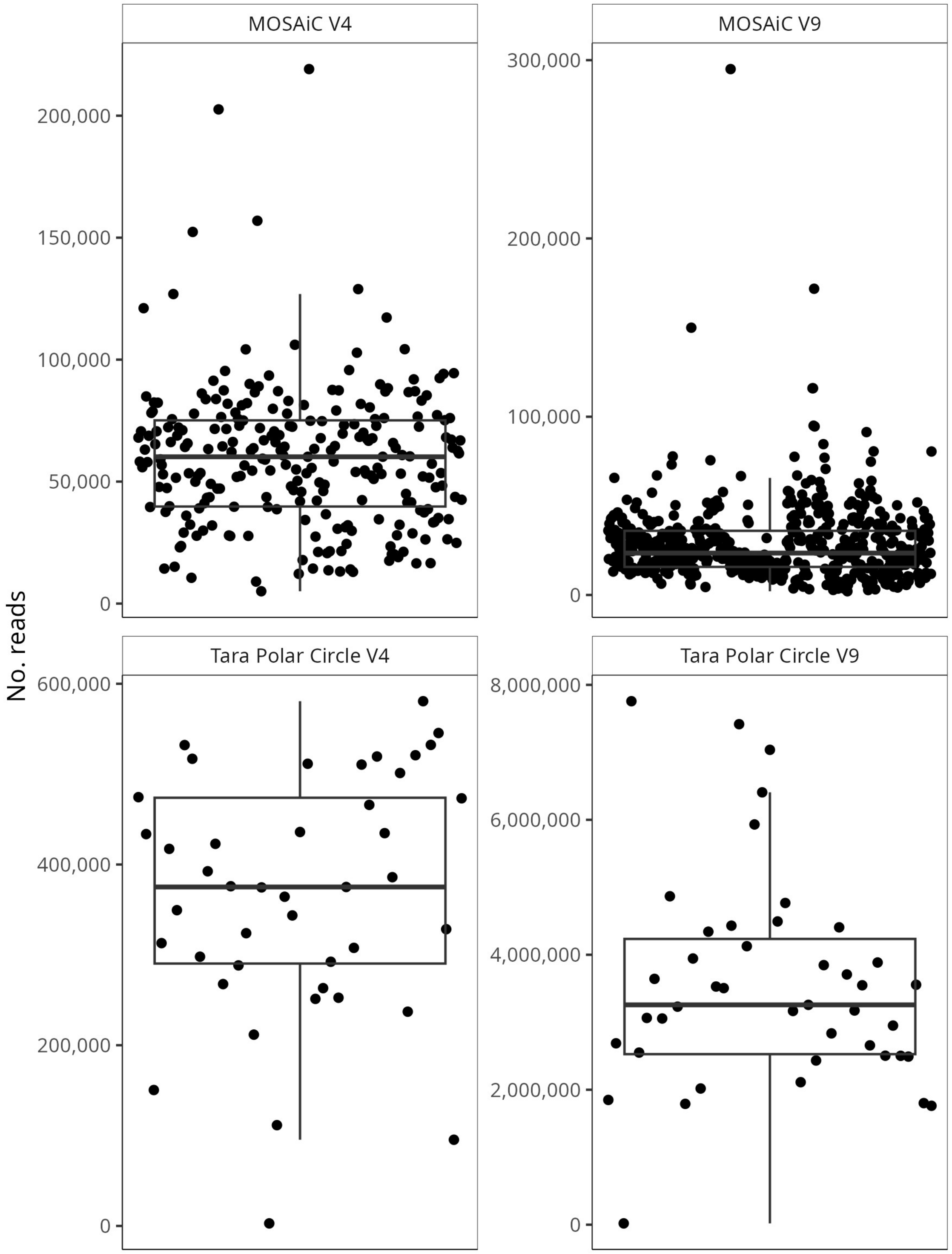


**Fig. S1:** Sequencing depth of the samples from the *Tara* Polar Circle and MOSAiC sampling campaigns after merging replicates and size fractions, and after selecting protist taxa. V4, V9: marker gene regions of the 18S rRNA gene.

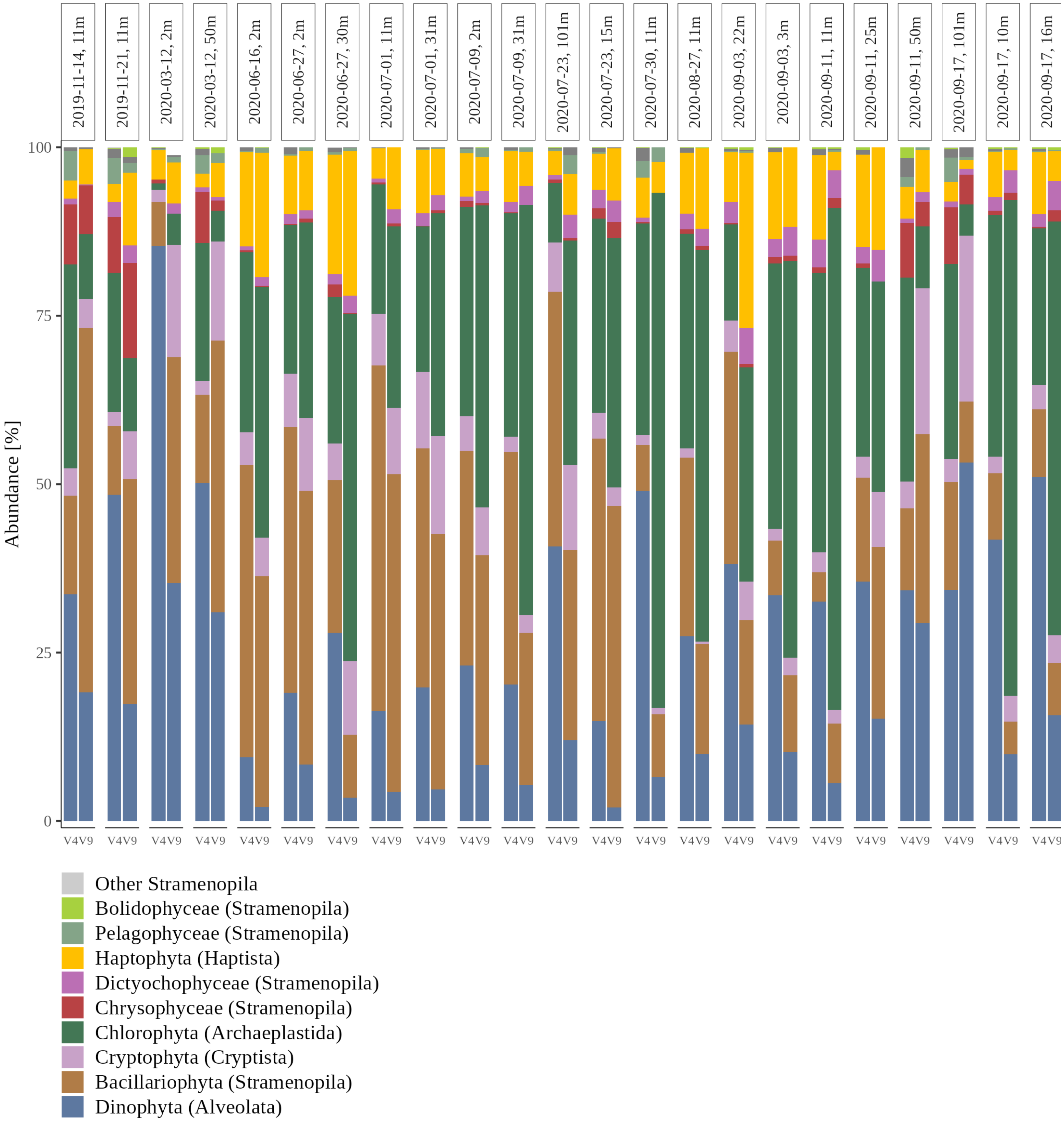


**Fig. S2:** 18S rRNA marker gene region comparison between V4 and V9. Shown are taxonomic assignments for all sampling points from which both markers were sequenced.

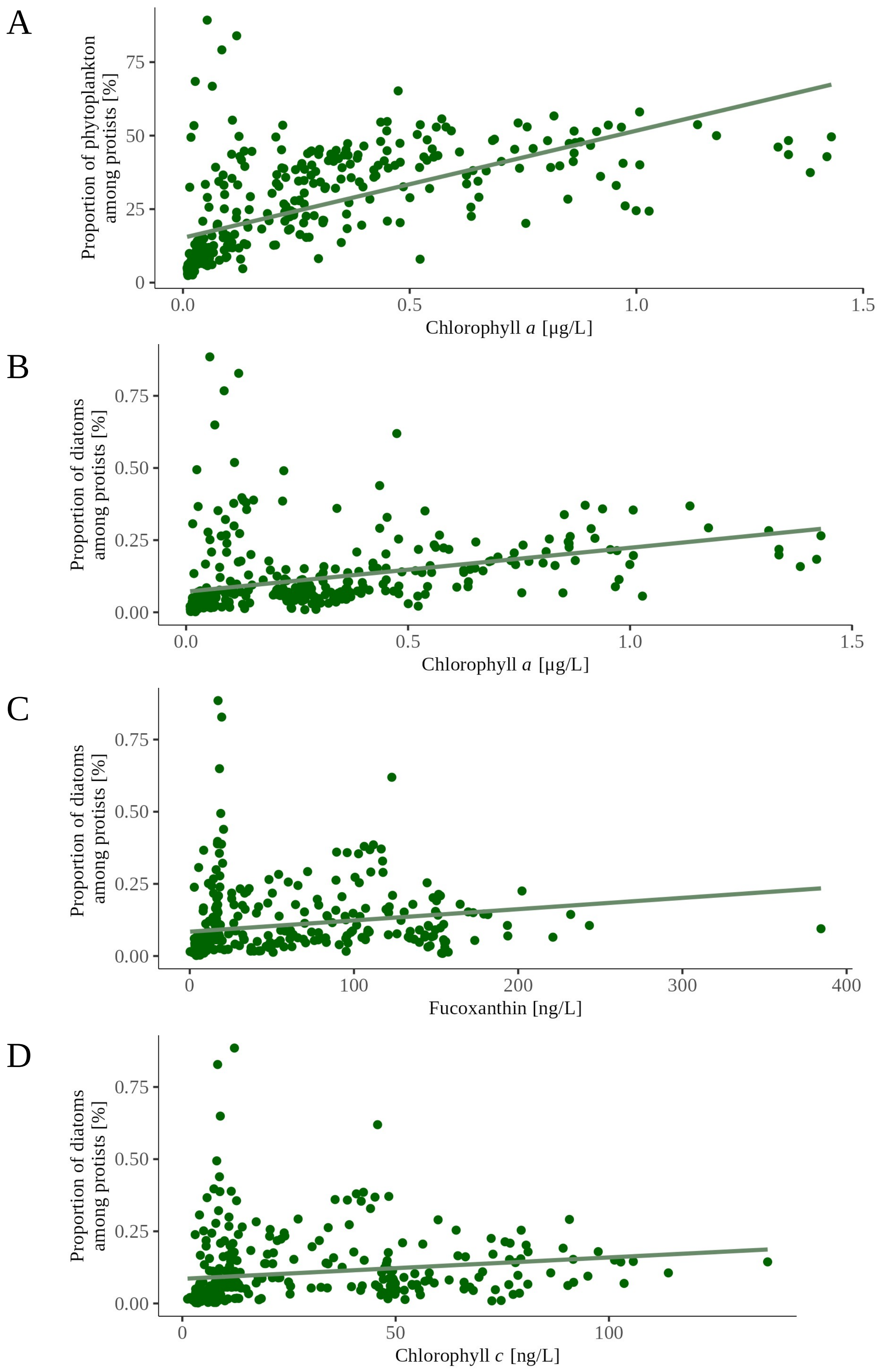


**Fig. S3:** Correlations between phytoplankton and pigment concentrations. (A) Proportion of phytoplankton among protists vs. chlorophyll *a*, Pearson's $r = 0.63$, $p < 0.005$, full dataset; (B) proportion of diatoms among protists vs. chlorophyll *a*, Pearson's $r = 0.36$, $p < 0.005$, full dataset; (C) proportion of diatoms among protists vs. fucoxanthin, Pearson's $r = 0.19$, $p < 0.005$, full dataset; (D) proportion of diatoms among protists vs. chlorophyll *c*, Pearson's $r = 0.17$, $p < 0.005$, full dataset.

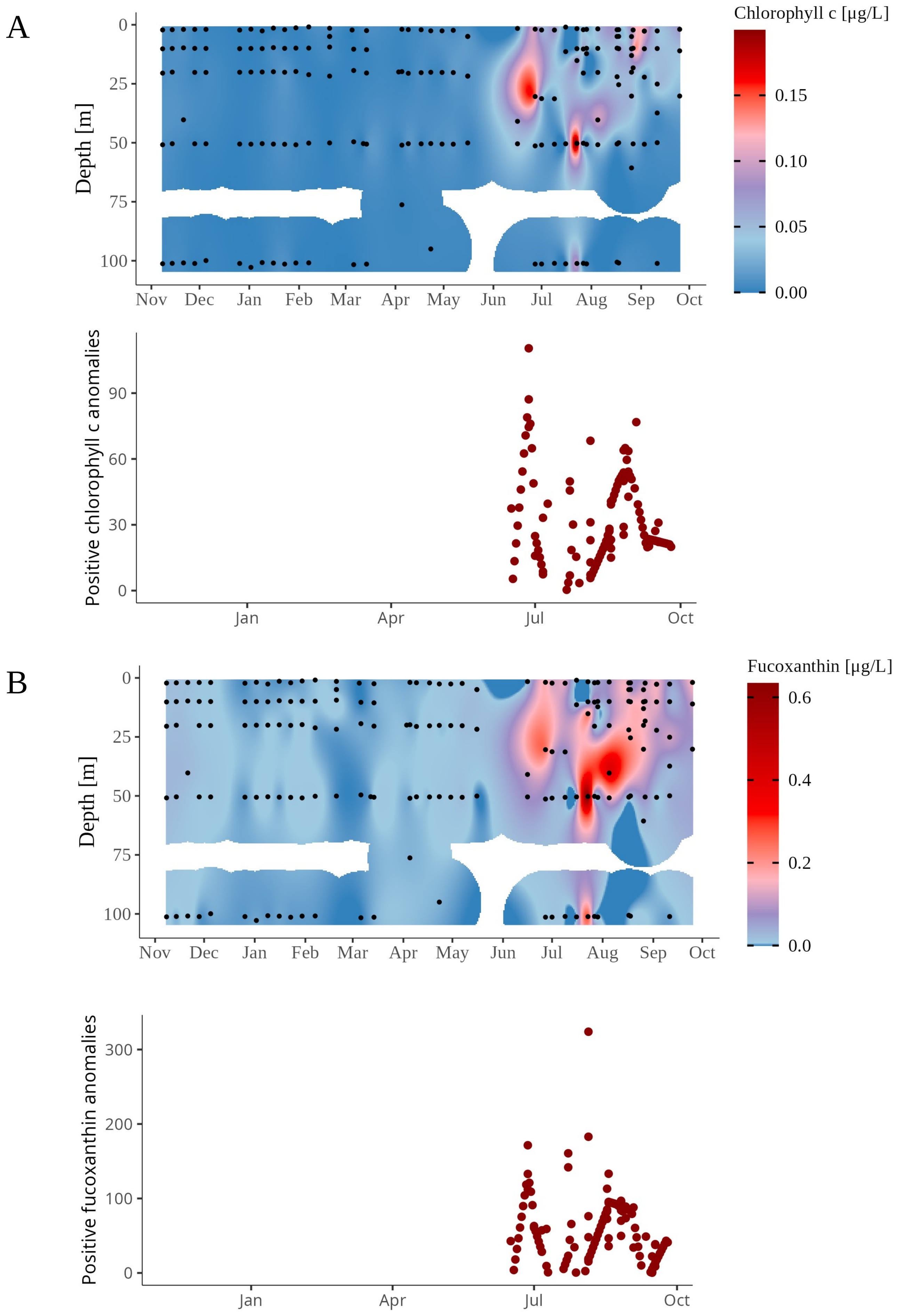


**Fig. S4:** (A) Chlorophyll *c* and (B) fucoxanthin pigment concentrations and anomalies (full dataset). Note the log-transformed scale.

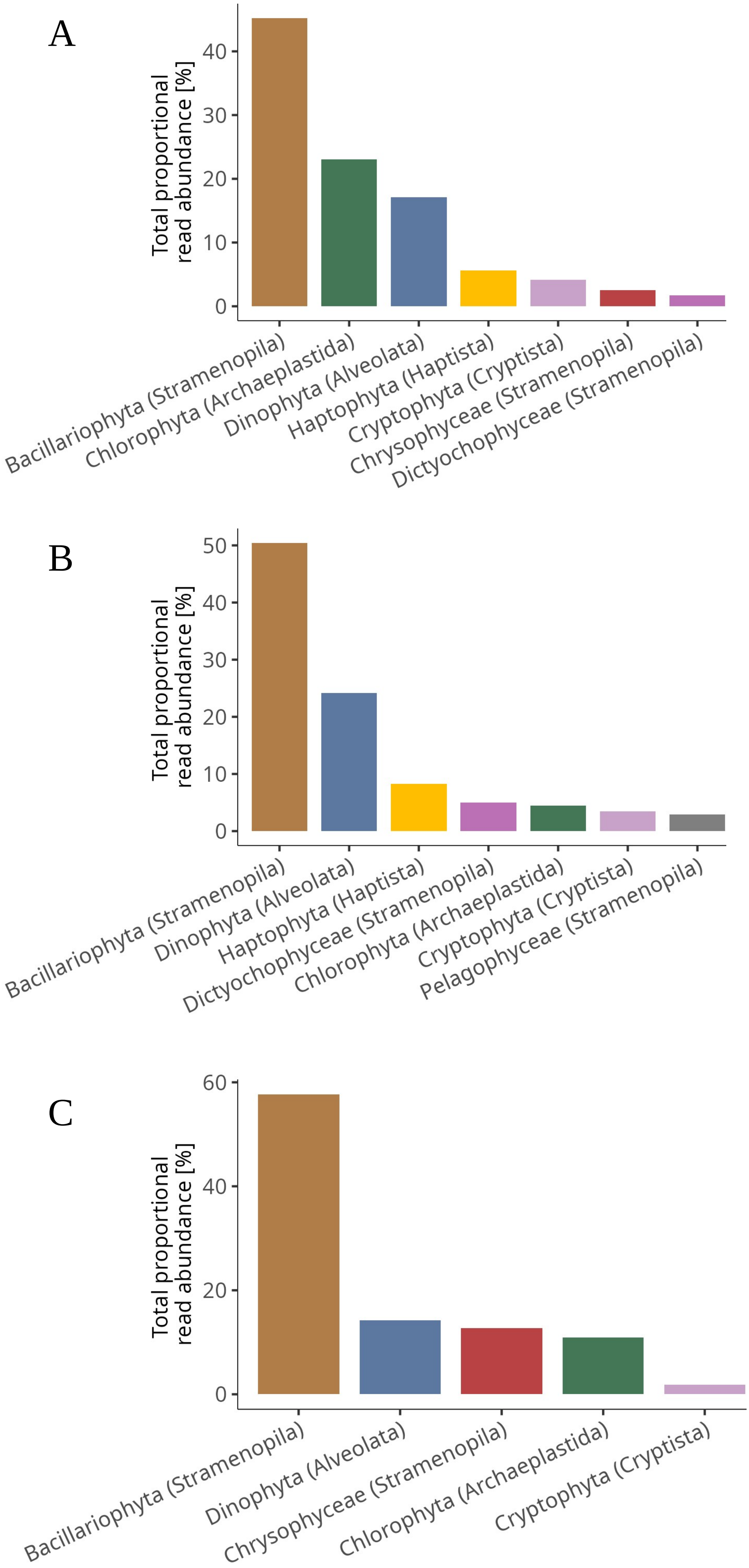


**Fig. S5:** Total proportional abundance of phytoplankton among protists per phytoplankton group in (A) under-ice water (time-series data); (B) ice-free water (time-series data); (C) sea ice. Only groups ≥1% are displayed.

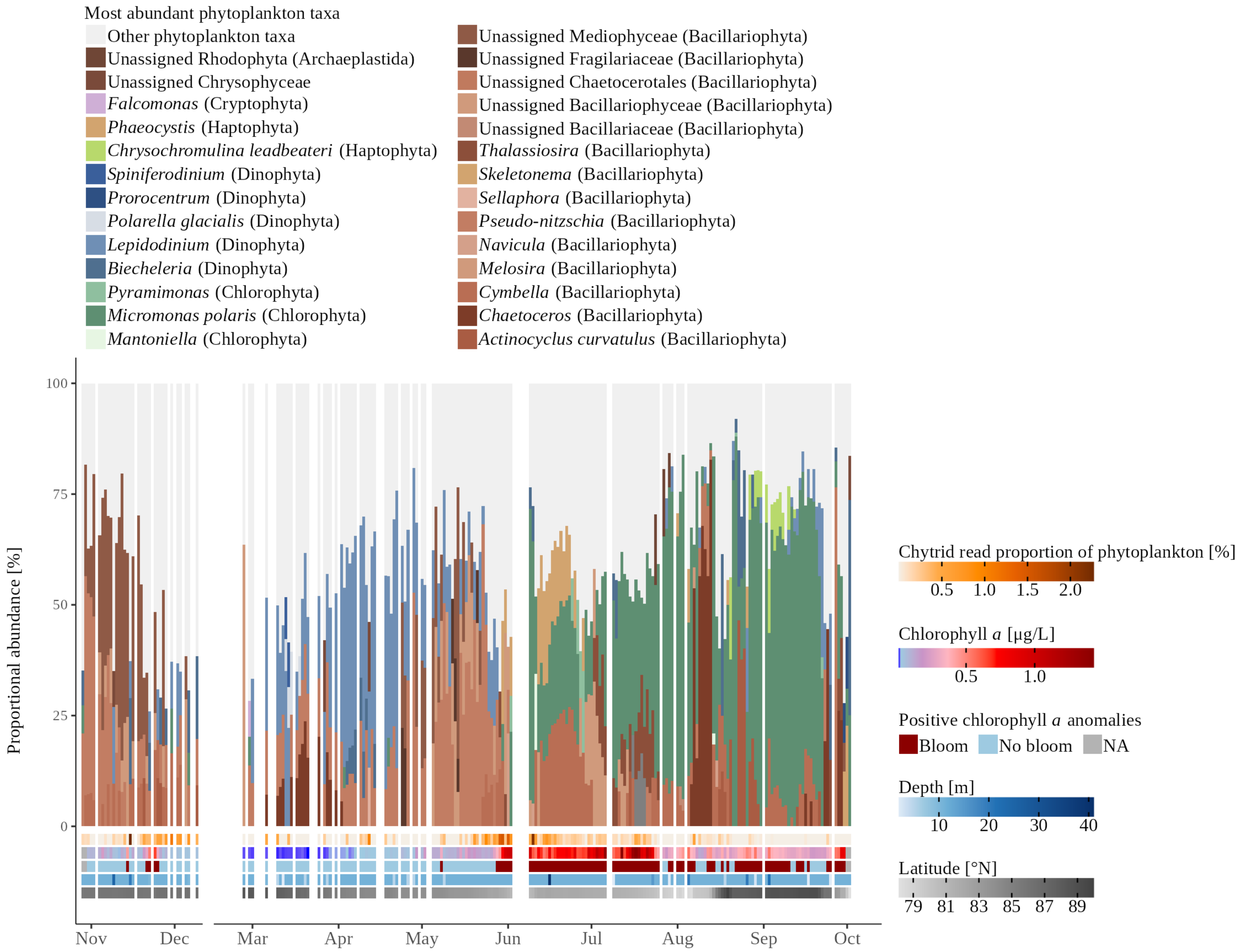


**Fig. S6:** The three most abundant ASVs among phytoplankton in each under-ice water sample as a proportion of phytoplankton (time-series data).

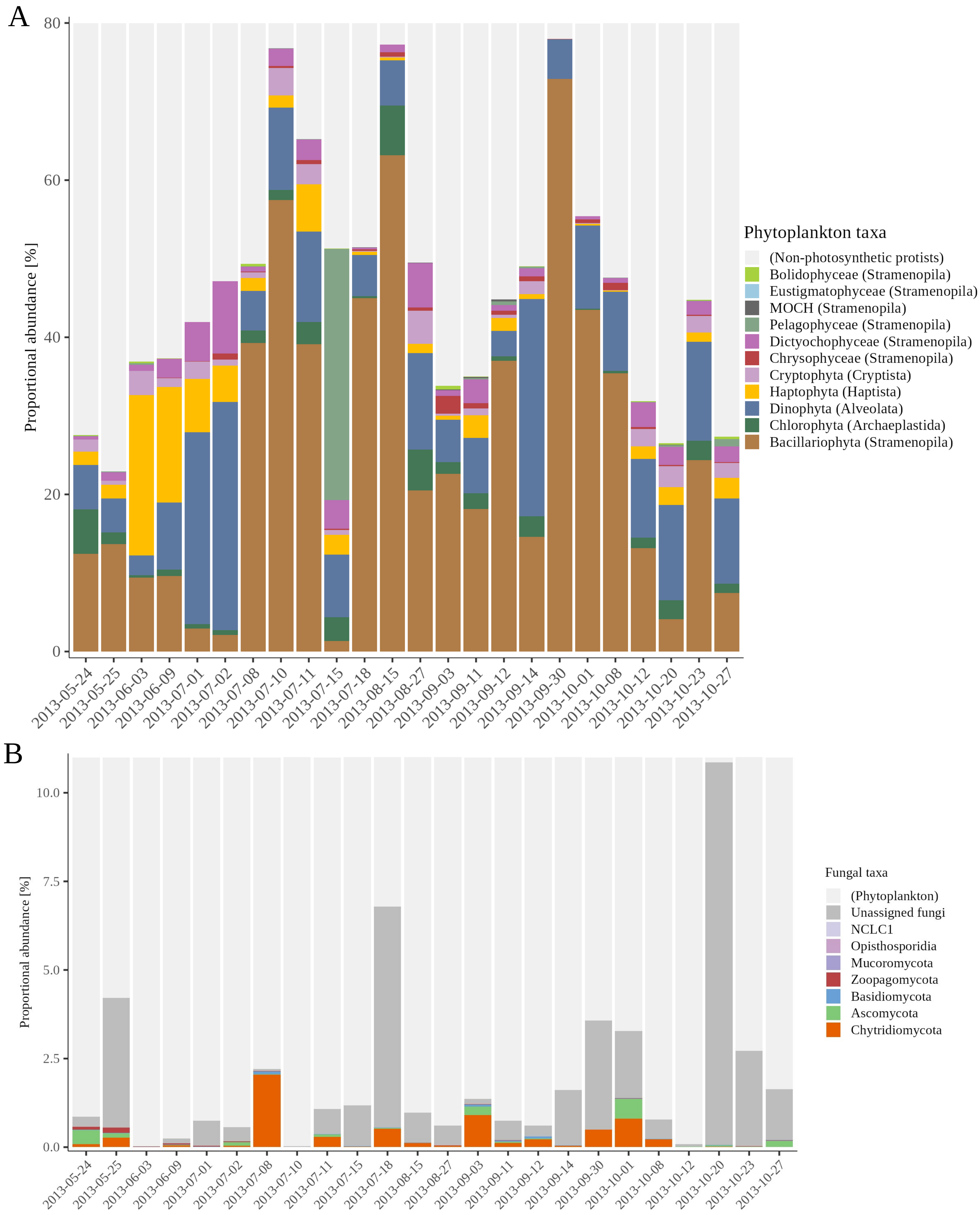


**Fig. S7:** Phytoplankton and mycoplankton communities in the ice-free water (time-series data). (A) Proportion of phytoplankton among protists; (B) proportion of fungi within the fungi + phytoplankton communities. NCLC1: Novel chytrid-like clade 1.

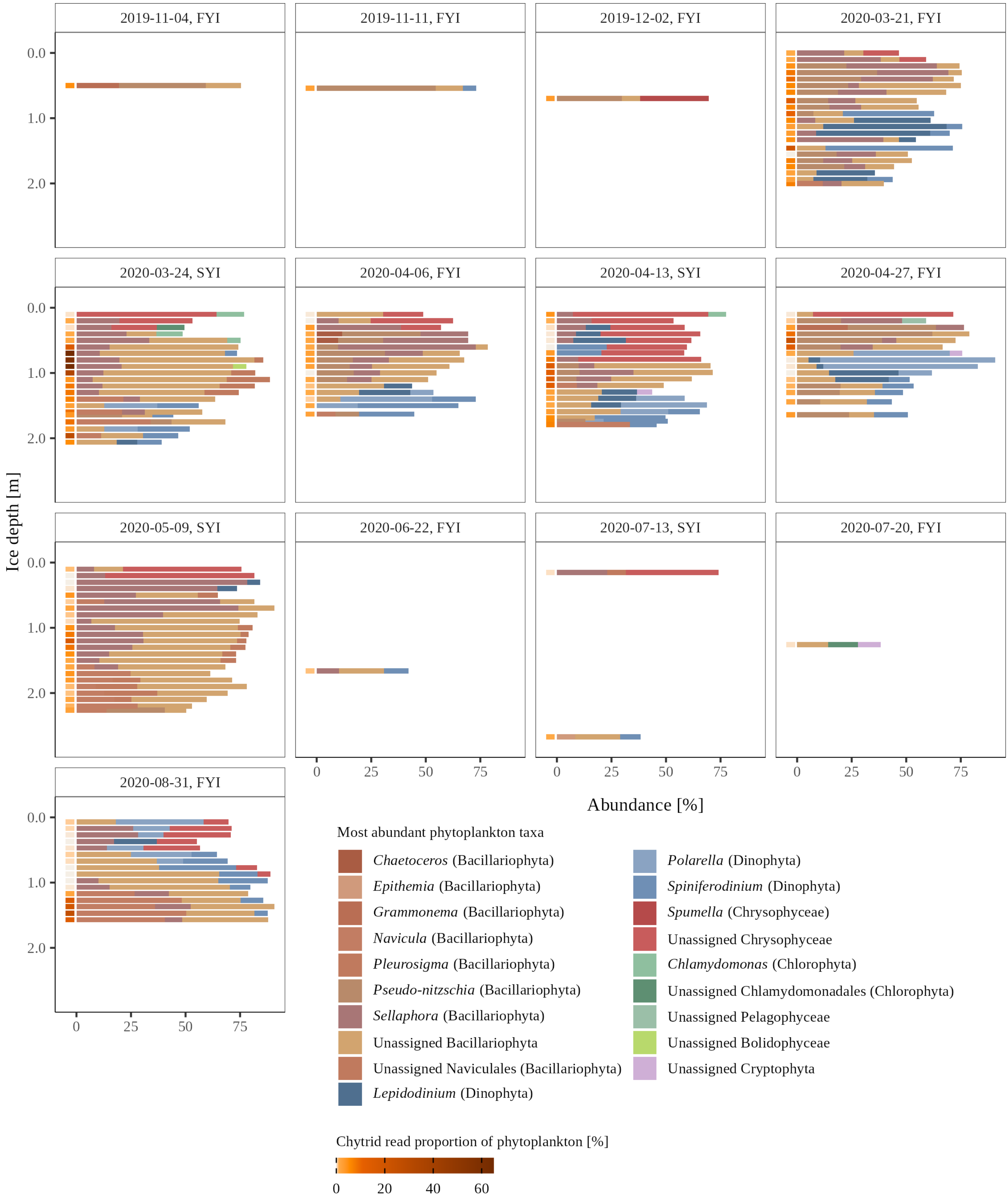


**Fig. S8:** Proportion of the three most abundant ASVs among phytoplankton in each sea-ice sample. FYI: first-year sea ice; SYI: second-year sea ice.

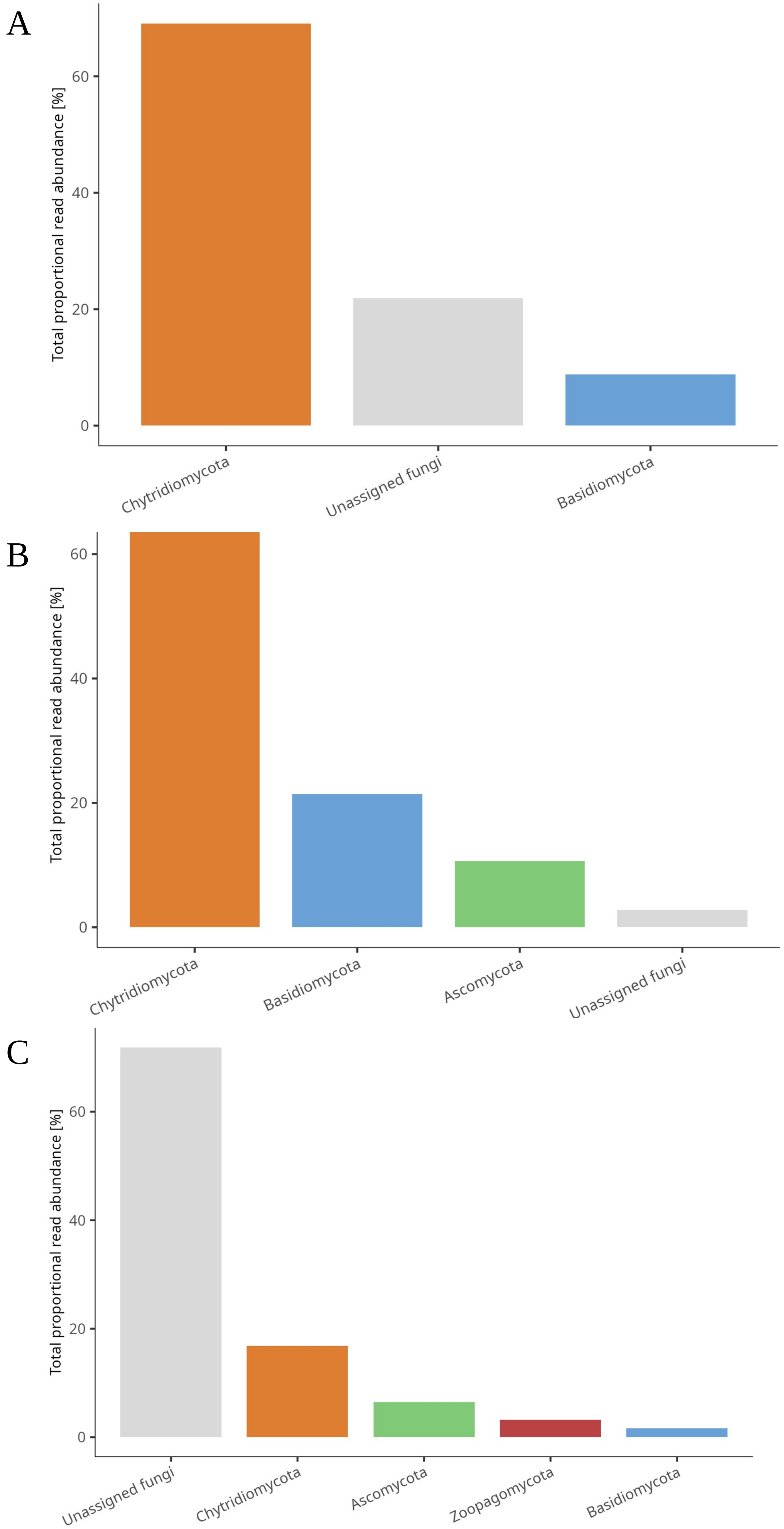


**Fig. S9:** Total proportional abundance of fungi among the fungi + phytoplankton communities per fungal phylum in (A) under-ice water (time-series data); (B) sea ice; (C) ice-free water (time-series data). Only groups representing ≥1% are displayed.

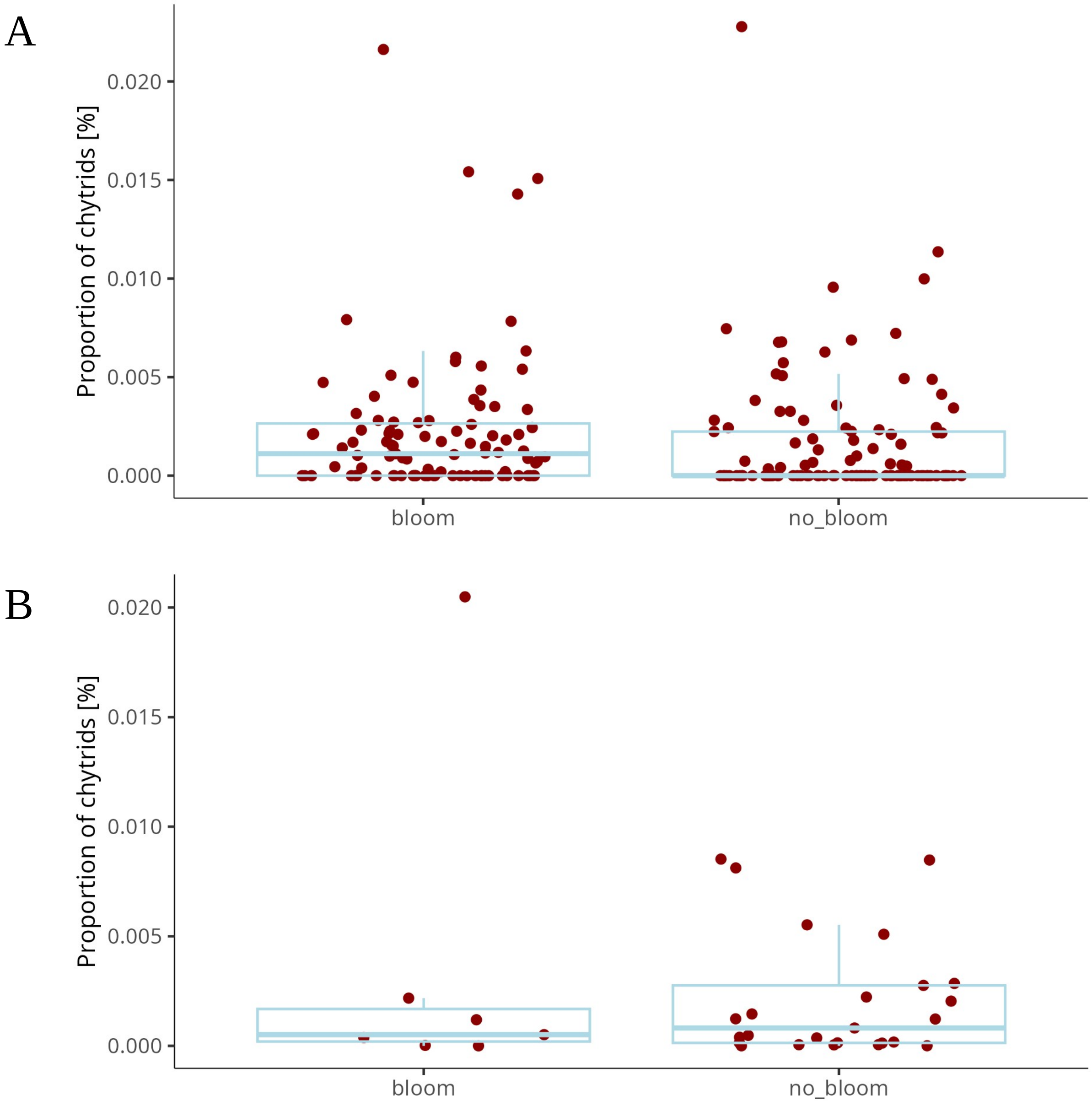


**Fig. S10:** Chytrid proportions of the chytrid + phytoplankton communities in bloom and non-bloom samples (threshold according to pigment anomalies; full dataset). Means were compared with the Wilcoxon test. (A) Under-ice water, $p < 0.001$; (B) ice-free water, $p = 0.72$.

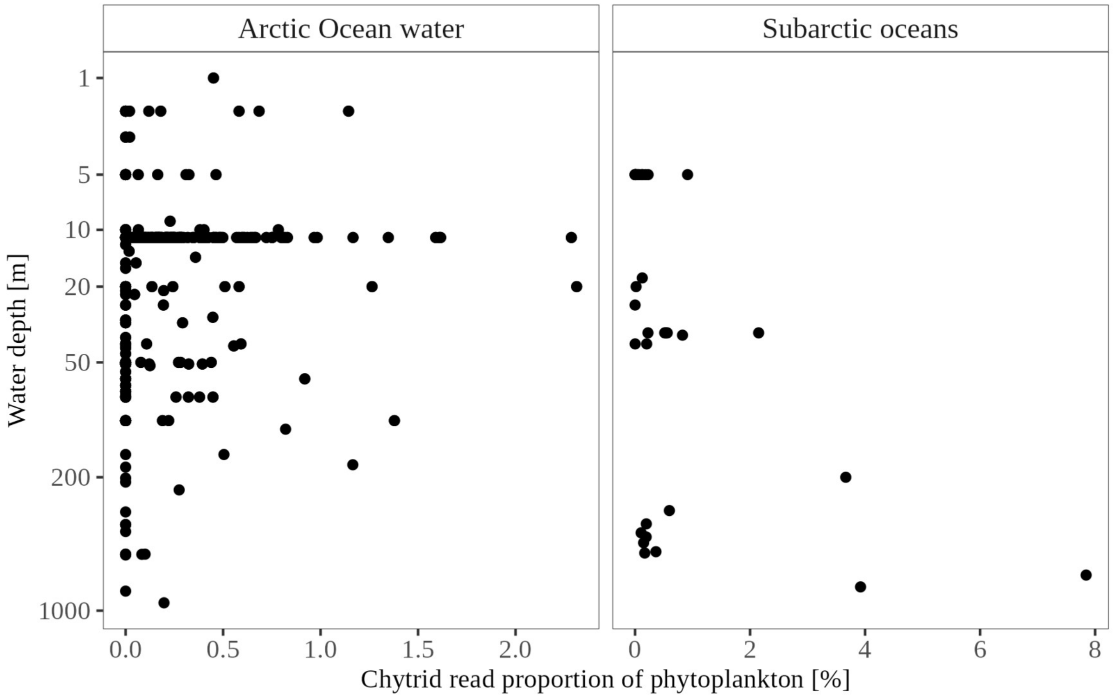


**Fig. S11:** Proportion of chytrids within the chytrid + phytoplankton communities per water depth. Note the log-transformed y-axes and different x-axes. Arctic Ocean water: under-ice water; Subarctic oceans: ice-free water.

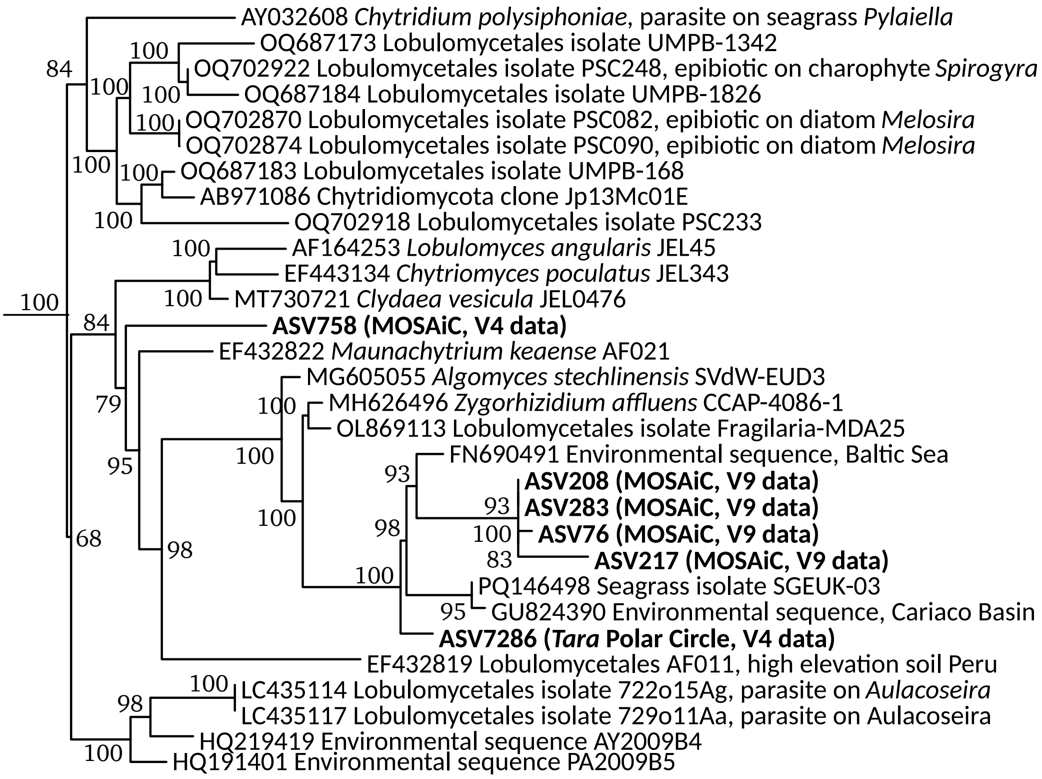


**Fig. S12:** Phylogenetic tree of the Lobulomycetales showing the affiliation of the most abundant chytrids from this study. Briefly, publicly available full SSU sequences of Chytridiomycota were aligned using MAFFT (Katoh and Standley, 2013, *Mol Biol Evol*) before new sequences were added with the --addfragments function. The alignment was filtered with trimAl (gap threshold 0.8; Capella-Gutiérrez et al., 2009, *Bioinformatics*) and a tree was inferred using IQ-TREE under the GTR+F+R9 model (Nguyen et al., 2015, *Mol Biol Evol*) with ultrafast bootstrap approximation (-bb 1000). Only the Lobulomycetales clade of the full tree is shown. Sequences from this study are in bold. V4, V9: marker regions of the 18S rRNA gene.

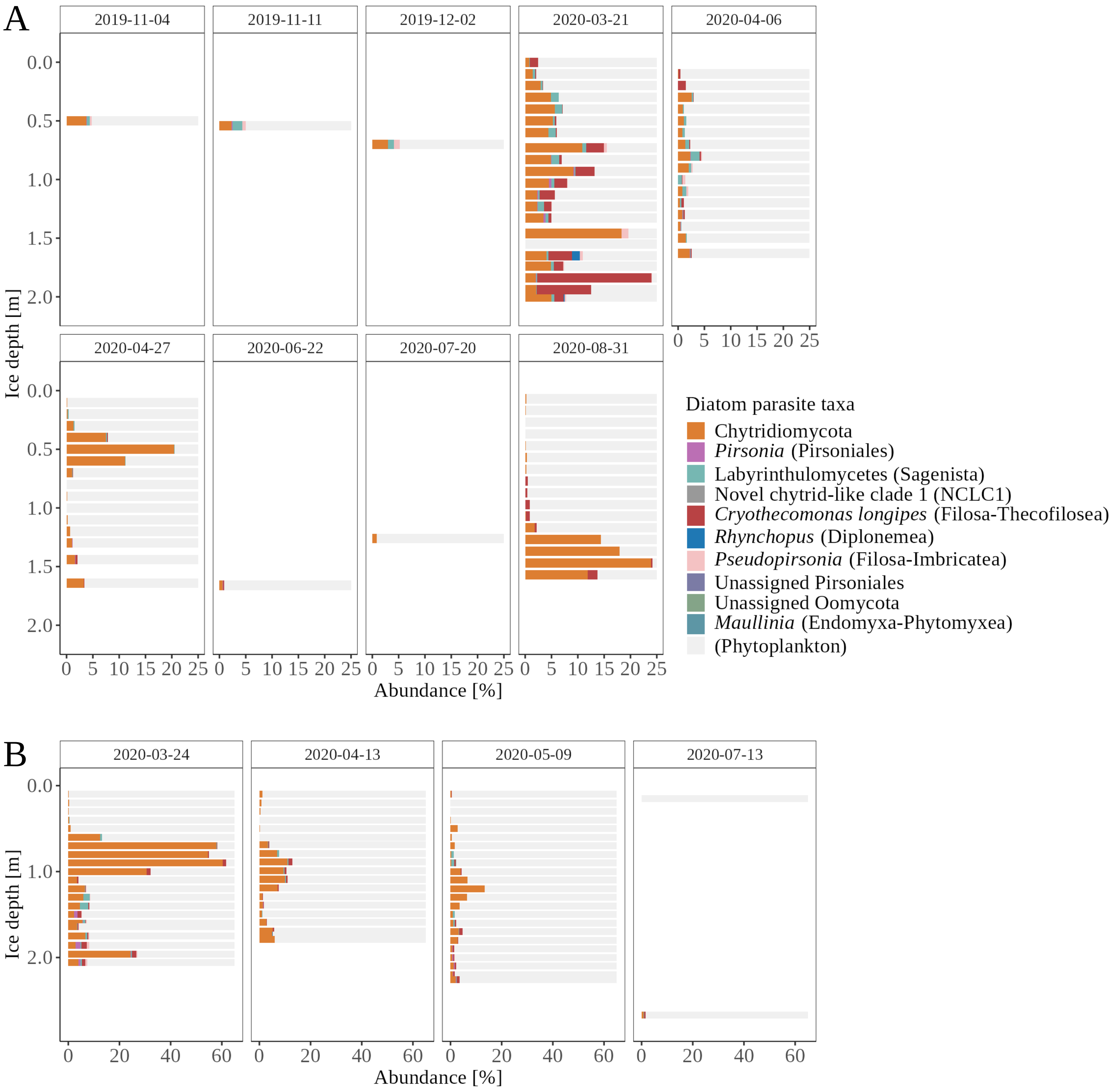


**Fig. S13:** Proportion of diatom parasites within the parasite + phytoplankton communities in the sea ice. (A) First-year sea ice; (B) second-year sea ice. White space: no data.

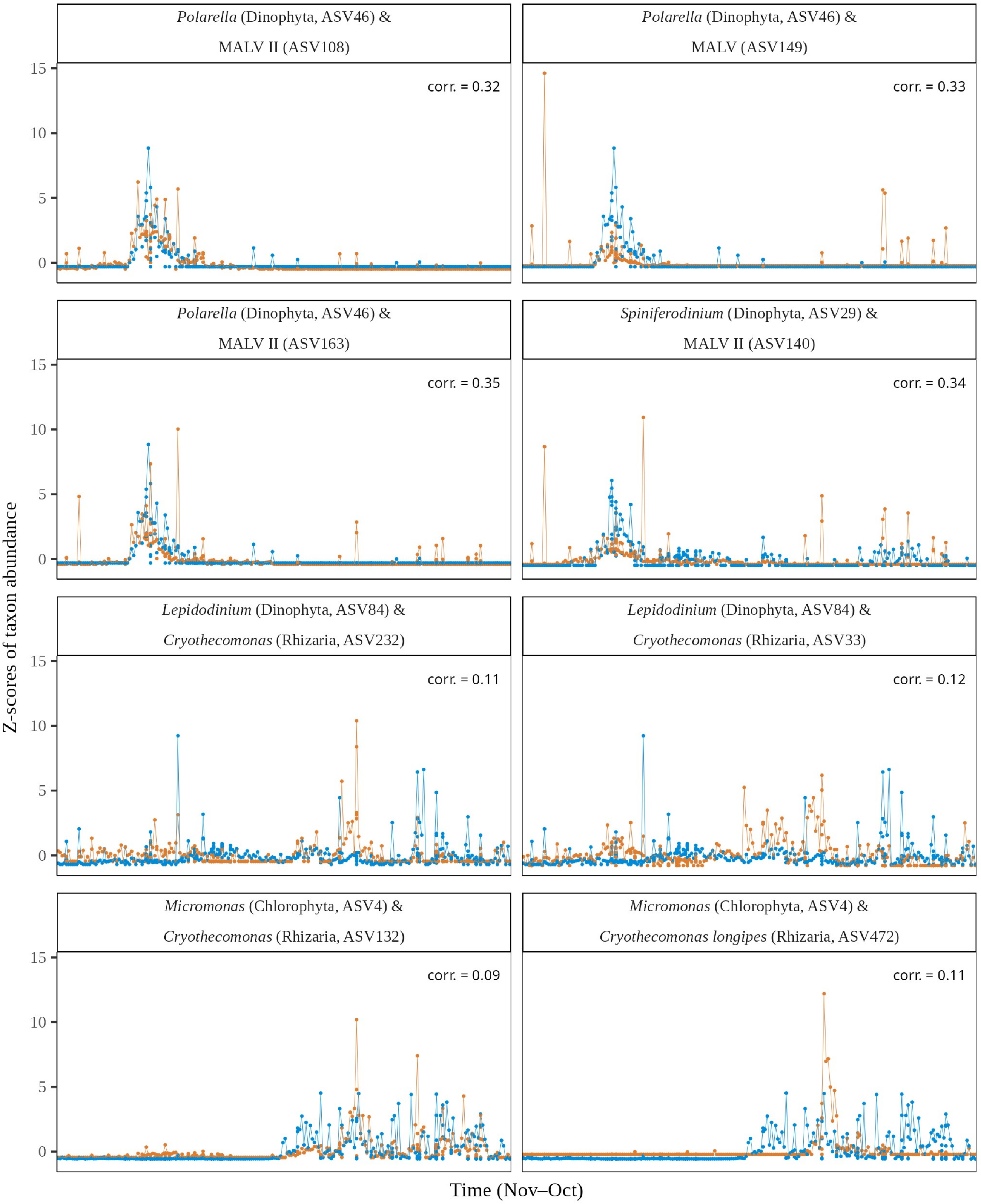


**Fig. S14:** *Z*-scores of raw read counts (taxon abundance) and significant Pearson correlation coefficients from Sparse Partial Least Squares regressions of selected non-chytrid parasite and phytoplankton ASVs within the parasite + phytoplankton communities in under-ice water, full dataset. Orange: parasite; blue: phytoplankton.

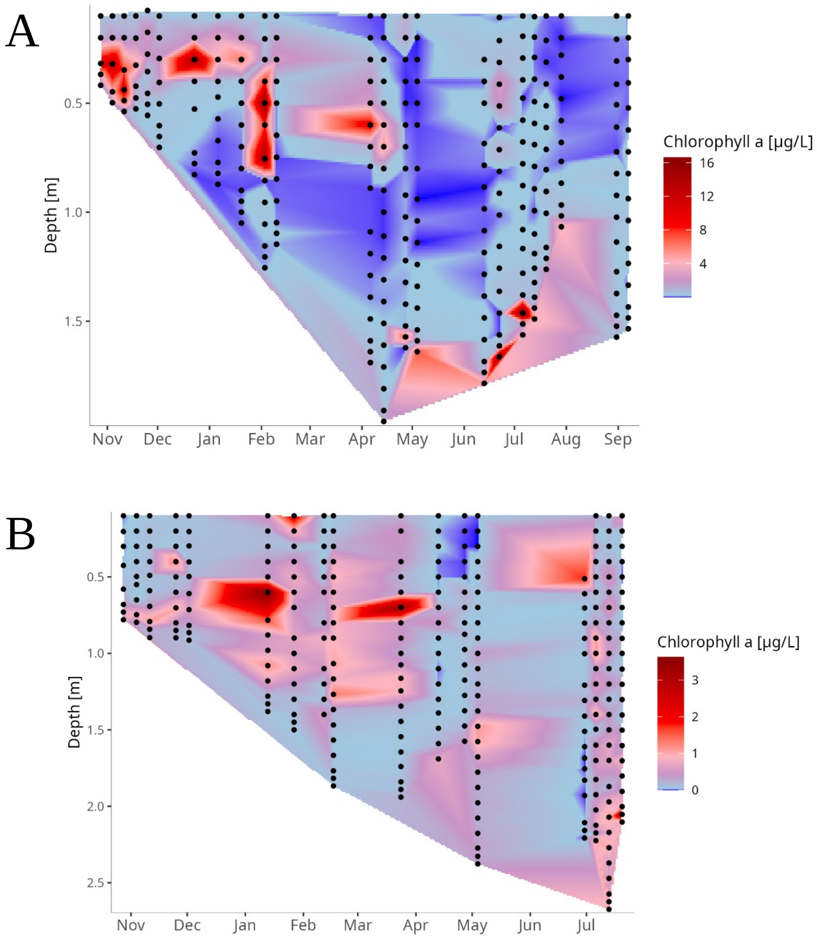


**Fig. S15:** Interpolated sea ice chlorophyll *a* concentrations. (A) First-year sea ice; (B) second-year sea ice. Black dots represent sampling points.

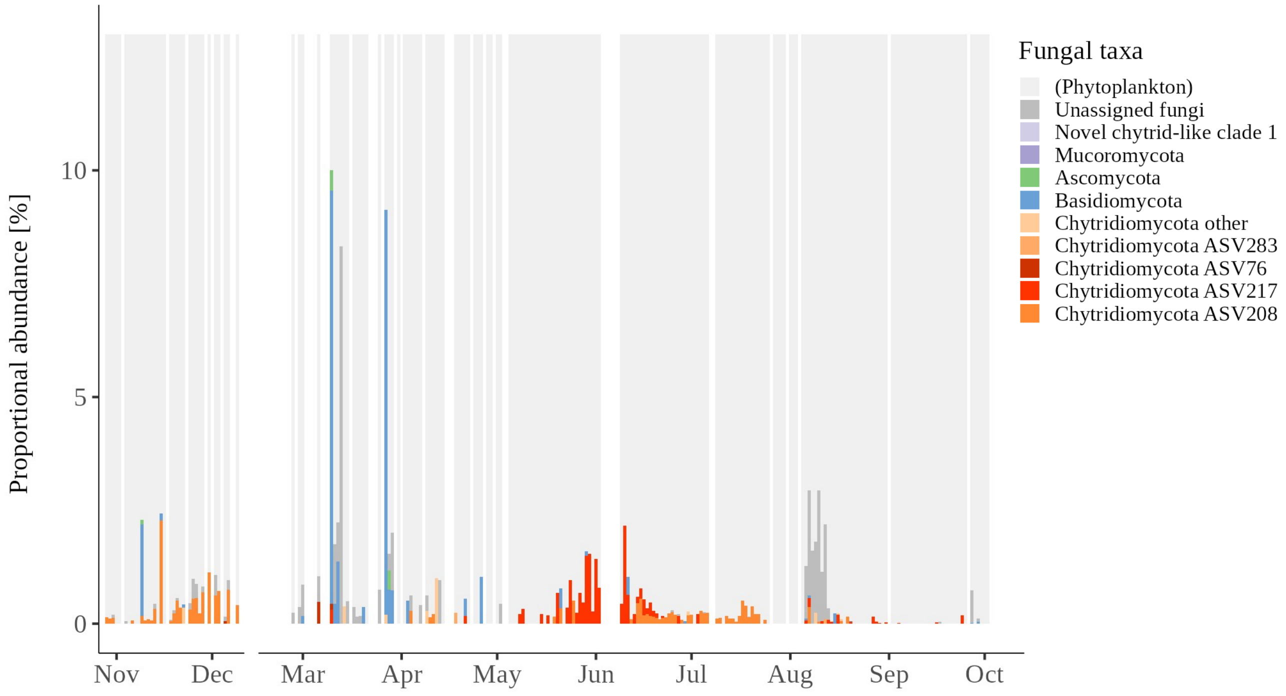


**Fig. S16:** Proportion of fungi within the fungi + phytoplankton communities in under-ice water, time-series data. This figure shows the full y-axis and complements Fig. 2C, where it is cropped for better visualisation of chytrids.
